# Temporal Dynamics of Faster Neo-Z Evolution in Butterflies

**DOI:** 10.1101/2023.12.19.572367

**Authors:** Lars Höök, Roger Vila, Christer Wiklund, Niclas Backström

**Affiliations:** Evolutionary Biology Program, Department of Ecology and Genetics (IEG), Uppsala University, Norbyvägen 18D, SE-752 36 Uppsala, Sweden; Institut de Biologia Evolutiva (CSIC-Univ. Pompeu Fabra), Passeig Martim de la Barceloneta 37-49, 08003 Barcelona, Spain; Department of Zoology: Division of Ecology, Stockholm University, Stockholm, Sweden

**Author notes:** Emails: Lars Höök: lars.hook[at]ebc.uu.se Roger Vila: roger.vila[at]csic.es Christer Wiklund: christer.wiklund[at]zoologi.su.se Niclas Backström: niclas.backstrom[at]ebc.uu.se.

**Keywords:** Faster-Z, Neo-sex chromosomes, Sex-biased gene expression, Lepidoptera, Selection

## Abstract

The faster-Z/X hypothesis predicts that sex-linked genes should diverge faster than autosomal genes. However, studies across different lineages have shown mixed support for this effect. So far, most analyses have focused on old and well differentiated sex chromosomes, but less is known about divergence of more recently acquired neo-sex chromosomes. In Lepidoptera (moths and butterflies), Z-autosome fusions are frequent, but the evolutionary dynamics of neo-Z chromosomes have not been explored in detail. Here, we analysed the faster-Z effect in *Leptidea sinapis*, a butterfly with three Z chromosomes. We show that the neo-Z chromosomes have been acquired stepwise, resulting in strata of differentiation and masculinization. While all Z chromosomes showed evidence of the faster-Z effect, selection for genes on the youngest neo-Z chromosome (Z3) appears to have been hampered by a largely intact, homologous neo-W chromosome. However, the intermediately aged neo-Z chromosome (Z2), which lacks W gametologs, showed less evolutionary constraints, resulting in particularly fast evolution. Our results therefore support that neo-sex chromosomes can constitute temporary hot-spots of adaptation and divergence. The underlying dynamics are likely causally linked to shifts in selective constraints, evolution of gene expression, and degeneration of W-linked gametologs which gradually expose Z-linked genes to selection.

## Introduction

Sex is often genetically determined by dimorphic sex chromosomes which initiate specific genetic pathways, resulting in the development of sex specific traits (Bachtrog et al., 2014). Sex chromosomes have evolved independently in different lineages from pairs of initially homologous autosomes (Wright et al., 2016). However, due to their unique inheritance pattern they eventually acquire properties that can lead to different evolutionary outcomes compared to the rest of the genome. The transition from autosomes to sex chromosomes starts with the acquisition of a sex-determining locus, followed by selection for reduced recombination between this locus and linked loci that benefit the same sex (Abbott et al., 2017). As a consequence, reduced recombination on the sex-limited chromosome generally leads to accumulation of deleterious substitutions, pseudogenization and eventually gene loss (Skaletsky et al., 2003; Wright et al., 2016). The degenerated state of the sex-specific chromosome together with the cessation of recombination can influence both the direction and intensity of selection on sex chromosomes (Charlesworth et al., 1987).

In female heterogametic species, genes on the Z chromosome are expected to evolve faster than autosomal genes, according to the faster-Z hypothesis (faster-X in male heterogametic systems) (Charlesworth et al., 1987) and multiple factors may contribute to this effect. In heterogametic females (ZW), recessive Z-linked mutations are immediately exposed to selection due to their hemizygous state, which can increase the rate of adaptive substitutions (Charlesworth et al., 1987). However, hemizygosity will also increase the efficiency of purifying selection, which can reduce interspecific divergence (Rousselle et al., 2016). Conversely, selection on dominant mutations which benefit males is expected to be efficient due to the male biased transmission of the Z chromosome (Charlesworth et al., 1987). In addition, both the lower copy number of Z chromosomes compared to autosomes and higher variance in male reproductive success will reduce the effective population size (*N_e_*) of Z-linked loci. This translates to a reduced efficiency of selection and a higher probability of fixation of slightly deleterious mutations on the Z chromosome, which can contribute to faster divergence compared to the autosomes (Vicoso & Charlesworth, 2009). Evidence for faster sex chromosome evolution have accumulated over the last decades, with examples from both male heterogametic organisms like *Drosophila* (Ávila et al., 2014), mice (Kousathanas et al., 2014) and humans (Veeramah et al., 2014), and female heterogametic snakes (Vicoso et al., 2013), birds (Mank, Nam, et al., 2010) and moths (Sackton et al., 2014). The pattern is, however, not universal (Rousselle et al., 2016) and the relative importance of selection and drift in driving sex chromosome evolution varies between species (Ávila et al., 2014; Wright et al., 2015). This variation shows that other factors than hemizygosity can influence rates of sex chromosome divergence (Wright et al., 2015). Gene content and expression patterns could also influence the faster-Z effect. For example, genes with sex-limited expression should have less selective constraints and could therefore be expected to evolve faster in general (Ellegren & Parsch, 2007). In addition, sex-linked genes are often specifically regulated through dosage compensating mechanisms (Gu & Walters, 2017). These mechanisms vary between species and can for example be achieved through hyperexpression or gene copy inactivation in one sex (Gu & Walters, 2017), which could affect the efficacy of selection for these genes (Mank, Vicoso, et al., 2010).

Faster-Z/X has predominantly been investigated in organisms with highly differentiated sex chromosomes and/or considerably degenerated sex-limited chromosomes (W/Y), but divergence of newly acquired neo-sex chromosomes is less explored. There are several properties that differ between ancestral and neo-sex chromosomes that need to be considered. First, functional gametologs (i.e. genes with homologs on the opposite sex chromosome), which should on average be more common on neo-sex chromosomes, can mask recessive mutations and thereby slow down divergence (Mrnjavac et al., 2023; Zhou & Bachtrog, 2012). This should reduce the adaptive potential of sex-linked genes in the heterogametic sex (Mrnjavac et al., 2023). Second, autosomal genes that become neo Z-linked will experience a new genomic environment which could lead to different evolutionary dynamics compared to genes that have been sex-linked for a long evolutionary time (Bachtrog et al., 2009; Mongue et al., 2022). Third, sex chromosome gene content and expression tend to get specialized and sex-biased over time (Mongue & Walters, 2018; Skaletsky et al., 2003). Early in this transition, however, there is likely a period of sexual conflict and pseudogenization events that will influence rates and directions of evolution (Nozawa et al., 2021). Taken together, the characterization of sequence and expression variation of neo Z-linked genes can give insights into the evolutionary processes that are no longer traceable on ancestral sex chromosomes.

In the order Lepidoptera (moths and butterflies), females are achiasmatic and lack meiotic recombination (Satomura et al., 2019; Turner & Sheppard, 1975). The presence of female achiasmy and the fact that a Z chromosome will spend 2/3 of the time in the male germline (assuming equal sex ratios) should increase recombination and thereby efficacy of selection on the Z chromosome relative to the autosomes (Hill & Robertson, 1966; Rousselle et al., 2019). Although estimates in butterflies suggest that other factors predominantly govern recombination rate variation between chromosome classes (Näsvall et al., 2023; Torres et al., 2022), female achiasmy needs to be considered in evolutionary studies of sex-chromosomes in butterflies. In Lepidoptera, the gene content of the Z chromosome is generally highly conserved across ∼ 250 MY of divergence (Wright et al., 2023). In addition, the W chromosome often lacks functional genes (Dalíková et al., 2017; Shipilina et al., 2022). Consequently, Z-linked genes have been hemizygous in females for most of lepidopteran evolution which makes it difficult to study how the early stages of sex chromosome evolution progressed. Fusions between autosomes and the ancestral Z chromosome have been frequent in Lepidoptera (Wright et al., 2023), but virtually nothing is known about how neo Z-linked genes evolve. A recent study showed that the neo-Z chromosome in the Monarch butterfly (*Danaus plexippus*) has evolved faster than the ancestral Z chromosome (Mongue et al., 2022), but the generality of this pattern and its mechanistic underpinnings require further study.

The wood white butterflies (genus *Leptidea*, Pieridae) have gained attention for their dynamic chromosome evolution, with multiple neo-sex chromosomes and extensively rearranged genomes (Šíchová et al., 2015; Yoshido et al., 2020; Höök, Näsvall et al., 2023). The observation that multiple sex chromosomes are present in several *Leptidea* species (Šíchová et al., 2016) suggests that their formations could predate the diversification of the genus ∼ 13 MYA (Wiemers et al., 2020). However, at least one of the neo-W chromosomes in the cryptic wood white species complex (*L. sinapis*, *L, reali* and *L. juvernica*) is less degraded and should therefore have become sex-linked more recently (Yoshido et al., 2020; Höök, Näsvall et al., 2023). The *Leptidea* system therefore provides a unique opportunity to study the early stages of sex chromosome evolution in butterflies. In addition, it opens the possibility to investigate how the presence of W-linked gametologs influences divergence of Z-linked genes.

Here we used previously available genome assemblies together with annotation and gene expression data for *Leptidea sinapis* and novel short-read sequencing data for four closely related species (*L. duponcheli, L. morsei, L. amurensis* and *L. lactea*) to characterize the temporal dynamics of sex chromosome evolution in detail. Our analyses show that the different sex chromosomes have been acquired in a step-wise fashion and that different stages of sex chromosome divergence are characterized by temporal variation in both selective constraints and levels of sex-biased expression.

## Methods

### DNA extraction and sequencing

DNA was extracted from the whole body of adult samples of two individuals of *L. duponcheli* (male + female), *L. morsei* (male + female), *L. amurensis* (males) and *L. lactea* (males), respectively, using the DNeasy kit (Qiagen). Libraries were constructed with Illumina TruSeq and 2×150 bp read length sequencing was done on a single Illumina NovaSeq6000 lane. Library preparations and sequencing were performed by NGI Stockholm (see acknowledgements).

### Read mapping

Raw sequence reads were trimmed for adapters and low quality bases with TrimGalore v.0.6.1 (Krueger, 2019) and Cutadapt v.4.0 (Martin, 2011). Contamination was assessed and filtered using FastQ Screen v.0.11.1 (Wingett & Andrews, 2018). Remaining singletons were removed using BBmap v.38.61b (Bushnell, 2019). Processed reads were mapped to the *L. sinapis* reference (Höök, Näsvall et al., 2023) with BWA mem v.0.7.17 (Li, 2013), discarding multi mapping reads. Duplicated reads were removed using MarkDuplicates from GATK v.4.1.1.0 (McKenna et al., 2010). Mapping statistics are presented in **Supplementary table 1**.

### Identification of Z-linkage

To identify Z-linked regions in the outgroups, we contrasted read depth between male and female samples for the two species (*L. morsei* and *L. duponcheli*) where data from both sexes were available. The male sample previously used to generate the *L. sinapis* assembly and an additional *L. sinapis* female sample was used as reference. Coordinates for *L. sinapis* genes were obtained from an available gene annotation and Z-linked genes were defined as ‘ancestral’ or ‘neo’ based on a previous synteny analysis (Höök, Näsvall et al., 2023). Read depth was calculated as the median of per position depth for each gene’s coding sequence (cds). In order to make comparisons between samples, the gene-wise medians were normalized by the mean of gene medians across autosomes. From the normalized values, gene-wise estimates for the Z chromosomes were calculated as log2 transformed male/female depth ratios. Genome-wide estimates were calculated as the mean of gene-wise estimates per chromosome. As a complement, we estimated SNP density across sex chromosome segments. In females, Z-linked genes without W-linked gametologs should lack SNPs while SNP density could be elevated for Z-W gene pairs if the W-linked copy has accumulated unique variants. Single sample variant calling was performed with bcftools mpileup v.1.17 (Li, 2011). The output was filtered to keep heterozygous sites with quality > 30. Depth (DP4) was limited to 25 - 200 and each allele was required to have a depth > 10. Filtered SNP counts were summarized in 100 kb windows, only including cds. To account for differences in average SNP counts, the window-based SNP densities were normalized by dividing the estimates for each sample by the median. For local estimates across the Z chromosomes, SNP densities were plotted in heatmaps, scaled by intraspecific maximum density of male Z-linked SNPs and square root transformed to improve visualization of low counts. Genome-wide SNP estimates were calculated as per chromosome mean of log2 transformed male/female SNP count ratios. Division by zero was allowed by adding 0.1 to each gene (depth) and window (SNPs). In addition, to assess the number of genes on neo-Z3 with putative W-linked gametologs, we used sequencing data from two additional *L. sinapis* samples (male and female). We defined Z-W gene pairs using two criteria: 1) a mean read depth in both female samples and the additional male control of > 80% of the read depth of the same gene in the male reference, and 2) the presence of SNPs in both female samples.

### Variant calling and estimation of population genetic summary statistics

To identify SNPs and estimate nucleotide diversity (ρε), we used 10 *L. sinapis* samples from an available resequencing data set (Talla et al., 2019). Variant calling and base quality score recalibration was performed using HaplotypeCaller from GATK v.4.1.1.0 (McKenna et al., 2010). The resulting all-sites vcf was filtered with bcftools v.1.14 (Li, 2011) using the cutoffs: -i ‘FMT/DP>5 & FMT/DP<25 & QUAL>30’. Coordinates for 0- and 4-fold degenerate sites, and synonymous and nonsynonymous SNPs (*P_n_* and *P_s_*) were identified with degenotate v1.2.0 (Sackton & Thomas, 2023). Window-based (100 kb sliding windows with 10 kb steps) estimates of ρε were calculated with pixy v.1.2.5.beta1 (Korunes & Samuk, 2021) and gene-wise estimates of Tajima’s D were generated with vcftools v.0.1.16 (Danecek et al., 2011).

### Estimation of divergence and selection

Gene coordinates and chromosomal assignment was based on available *L. sinapis* annotation and synteny information (Höök et al., 2023). For the outgroup species, cds were generated by making consensus sequences from the read mapping performed against the *L. sinapis* reference, using Samtools v.1.16.1 (Li et al., 2009), based on the *L. sinapis* gene coordinates. Prior to alignment, all genes were scanned for intact open reading frames (ORF) with Emboss getorf v.6.6.0 (Rice et al., 2000), selecting the longest ORF for each gene. Codon aware alignments between all species were made with Prank v.170427 (Löytynoja & Goldman, 2005). Poorly aligned regions were filtered using HmmCleaner v.0.180750 (Di Franco et al., 2019) and only sites represented in more than three species after filtering were retained using reportMaskAA2NT from MACSE v.2.06 (Ranwez et al., 2011). Alignments with < 50% of their initial length were discarded. In total, filtering removed 20.38% of autosomal and 22.11% of Z-linked genes which resulted in 9,906 autosomal-, 511 ancestral Z-, and 997 neo Z-linked genes. Interspecific divergence was estimated using gene-wise *d_N_* / *d_S_*-ratios (μ) with the branch model (NSsites = 0, model = 2) in Paml v.4.9j (Yang, 2007). Since substitutions can be confounded by polymorphisms, we masked SNPs in the *L. sinapis* reference before estimating divergence. Genes with μ = 999 (lack of synonymous substitutions) were filtered out (autosomal: 5.64%, ancestral Z: 7.83%, and neo-Z: 8.22%). In addition, to account for genes without synonymous substitutions and to normalize for gene size, we separately calculated point estimates of μ as the ratio of the sum of non-synonymous substitutions (*D_n_*) divided by the sum of non-synonymous sites (*N*), and the sum of synonymous substitutions (*D_s_*) divided by the sum of synonymous sites (*S*) across all genes per chromosome category (*D_n_* / *N* / *D_s_* / *S*) (Mank *et al*., 2007). Direction of selection (DoS) was calculated per gene as *D_n_* / (*D_n_* + *D_s_*) -*P_n_* / (*P_n_* + *P_s_*), where *P_n_* and *P_s_* are the number of non-synonymous and synonymous polymorphisms (Stoletzki & Eyre-Walker, 2011). Codon usage bias was estimated with coRdon v.1.16.0 (Elek et al., 2023) using the ENC’ function. GC content at 4-fold degenerate sites was calculated with a custom script.

### Gene expression quantification

Gene expression data for *L. sinapis* were extracted from previous studies (Leal et al., 2018; Höök *et al*., 2019). Briefly, raw RNA-seq reads were trimmed with the same methods used for the DNA reads (see Read mapping) with additional trimming using prinseq v.0.20.4 (Schmieder & Edwards, 2011) to remove poly A/T/N tails, low complexity regions and reads with > 10% N’s. Trimmed reads were aligned to the reference assembly of *L. sinapis* (Höök, Näsvall et al., 2023) with STAR v.2.7.9.a (Dobin et al., 2013) guided by gene annotation. Expression levels were quantified using Stringtie v.2.1.4 (Pertea et al., 2015) and sex-biased expression was inferred with Deseq2 v.1.20.0 (Love et al., 2014).

### Statistical analyses

Statistical analyses were performed in R v.4.2.3 (R Core Team, 2023) and figures were generated with ggplot2 v.3.4.2 (Wickham, 2009). For multiple comparisons, we performed Kruskal-Wallis rank sum tests followed by pairwise Wilcoxon rank sum tests with Bonferroni correction. Bootstrap statistics were performed with 10,000 replications. Permutation tests were performed to assess the difference in μ between chromosomal categories. Chromosome assignment was randomized and for each permutation the difference in μ was calculated to generate a null distribution to which the empirical difference was compared. Two-tailed p-values were calculated as the number of times a more extreme value than the empirical estimate was observed in the null distribution, corrected with the Bonferroni method. To test for a faster-Z effect, including the effects of GC% at 4-fold degenerate sites and gene expression levels, linear modelling was performed with the lm function in R using the formula: log(μ) ∼ chromosome type + 4-fold GC% + log(FPKM). Genes without non-synonymous substitutions were removed to normalise the residuals. Model assumptions were assessed with diagnostic plots and a non-constant variance score test (*χ^2^* = 0.488, df = 1, p = 0.485).

## Results

### Stepwise acquisition of neo-Z chromosomes

The Z chromosomes in *Leptidea sinapis* and the closely related *L. reali* and *L. juvernica* have recently been characterized (Yoshido et al. 2020; Höök, Näsvall et al. 2023), but it is not known if the same genomic regions are Z-linked in other *Leptidea* species. We therefore inferred Z-linkage for two additional species: *L. morsei*, representing the sister clade to the three previously studied species, and *L. duponcheli*, representing the deepest split in the genus (**Figure 1**). The largest Z chromosome (Z1), which contains the ancestral lepidopteran Z chromosome fused with an autosomal part (homologous to *Bombyx mori* autosome 17), was generally male-biased for both read depth and SNP density in all *Leptidea* species. This shows that this fusion occurred before the radiation of all *Leptidea* species ∼13 MYA (Wiemers et al., 2020) (**Figure 1**). The second Z chromosome (Z2) was also male-biased in *L. morsei*, but in *L. duponcheli* the region corresponding to a part of *B. mori* autosome 11 was non-biased, implying it became Z-linked after the split between *L. duponcheli* and the rest of the clade (**Figure 1**). Finally, the smallest Z chromosome (Z3), did not show any bias in either *L. morsei* or *L. duponcheli* except for two female biased read depth peaks, largely overlapping in the different species (**Figure 1**). These peaks likely represent several genes occurring in multiple copies on the W chromosomes in the different species. It should be noted that the identified Z-linked regions in *L. duponcheli* and *L. morsei* most likely occur in different configurations compared to *L. sinapis*, since the Z chromosomes are rearranged even between the closely related *L. sinapis*, *L. reali* and *L. juvernica*. In *L. sinapis,* several regions had elevated SNP density compared to the male (**Figure 1**), which is expected if Z/W gametologs have retained homology, but also accumulated unique variants. SNP density ratios for individual chromosomes clearly showed that Z3 differs from the autosomes in *L. sinapis,* with an increased SNP density in the female (**Supplementary figure 1**). In contrast, for *L. duponcheli* and *L. morsei*, Z3 did not deviate from the autosomes (**Supplementary figure 1**). This points to a recent acquisition of Z3, i.e. after the split between *L. morsei* and the sister taxa *L. sinapis*, *L. reali* and *L. juvernica* ∼3-4 MYA (**Figure 1**). To assess the number of genes on Z3 with preserved W gametologs, we used additional sequencing data from two samples of *L. sinapis* (Catalan population, male and female) mapped against the *L. sinapis* reference. Of the 300 genes included in the gene set, 251 had comparable or higher read depth than the male reference in all samples while 24 genes had reduced depth (< 80%) exclusively in the two female samples. We found 279 genes with SNPs in both female samples and 12 genes where SNPs were not found in either of the female samples. The intersect of genes with both unbiased read depth and female SNPs was 244, suggesting that a vast majority (∼80%) of genes on Z3 have retained W gametologs (**Supplementary figure 2**).

**Figure 1.**
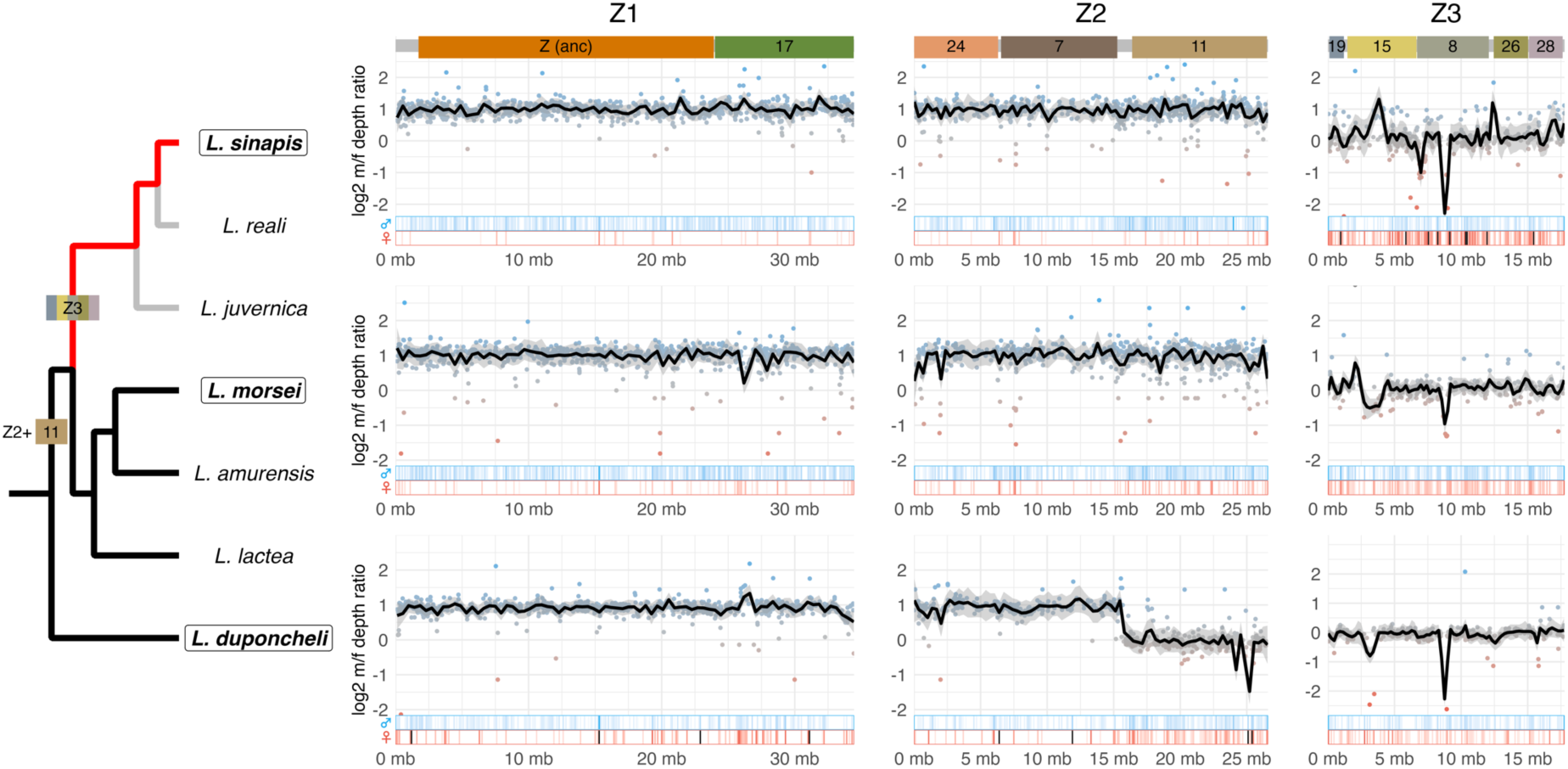
Read depth ratios and SNP densities in males and females based on reads mapped to the male L. sinapis Z chromosomes. The top panel shows syntenic autosomal regions in B. mori (Höök, Näsvall et al. 2023) and the cladogram shows the inferred events leading to formation of neo-Z chromosomes Z2 and Z3. The cladogram is based on (Dincă et al., 2011). The focal branch for the analysis of divergence is highlighted in red (L. sinapis) and the outgroup branches are in black (L. morsei, L. amurensis, L. lactea and L. duponcheli). Grey lines indicate species not included in the analysis (L. reali and L. juvernica). Species for which the read depth and SNP density analysis was performed are highlighted in bold and circled (L. sinapis, L. morsei and L. duponcheli). Dot plots show gene-wise depth ratios with a loess line, smoothed over 1 Mb. Heatmaps show the male and female SNP densities in 100 kb windows, scaled by the maximum estimates of the male Z chromosomes (values above max are highlighted in black) and square root transformed to improve visualization. Data for all chromosomes are summarized in **Supplementary figure 1**.

Although several shorter autosomal regions had deviating mapping ratios, which could represent translocations of individual genes, the general patterns suggest that no additional autosomes have been fused to the sex chromosomes in the outgroup species (**Supplementary figure 3**).

### Faster evolution on both the ancestral and the neo-Z chromosomes

After characterizing the temporal acquisition of sex chromosomes in the *Leptidea* system, we investigated the evolutionary rates for genes on the different Z chromosomes and the autosomes. We found an elevated μ for genes on both ancestral- and neo-Z chromosomes compared to autosomal genes (Kruskal-Wallis test, *χ^2^* = 84.408, p < 2.20*10^-16^; Wilcoxon tests, p < 2.10*10^-5^; **Table 1**, **Supplementary table 2**). For the neo-Z chromosomes, both *d_N_* and *d_S_* were significantly higher than for the autosomes (Wilcoxon test, p < 1.80*10^-8^; **Supplementary table 2**), while the higher μ for the ancestral Z chromosome was a result of both a significantly higher *d_N_* (Wilcoxon test, p = 4.84*10^-3^; **Supplementary table 2**) and significantly lower *d_S_* (Wilcoxon test, p = 2.00*10^-2^; **Supplementary table 2**, **Figure 2A**). Note that both *d_N_* and *d_S_* (but not μ) were significantly different between the ancestral and the neo-Z chromosomes (Wilcoxon test, p < 6.30*10^-4^; **Supplementary table 2**, **Figure 2a**). The reduced *d_S_* on ancestral Z was not caused by codon usage bias, which was significantly reduced on the ancestral Z compared to the other chromosome categories (median ENC’: A = 58.4, ancestral Z = 59.4, neo-Z = 58.3; Kruskal-Wallis test, *χ^2^* = 27.534, p = 1.05*10^-6^; Wilcoxon tests, p < **6.42*10^-5^**, **Supplementary table 3**). However, there were no significant differences in estimates of divergence between the ancestral and the neo-Z part of Z1 (Wilcoxon tests, p > 0.113; **Supplementary table 2**, **Figure 2B**). As an additional measure, we estimated μ based on the sum of all substitutions across each chromosome class, thereby including genes without any synonymous substitutions. Again, we found significantly higher μ for Z-linked (median ancestral Z = 0.1868, neo-Z = 0.1780) compared to autosomal genes (median = 0.1278) (Permutation test, 10,000 repetitions, p < 6*10^-4^), but no difference between genes on the neo and the ancestral Z (Permutation test, p = 1; **Supplementary figure 4**).

**Figure 2.**
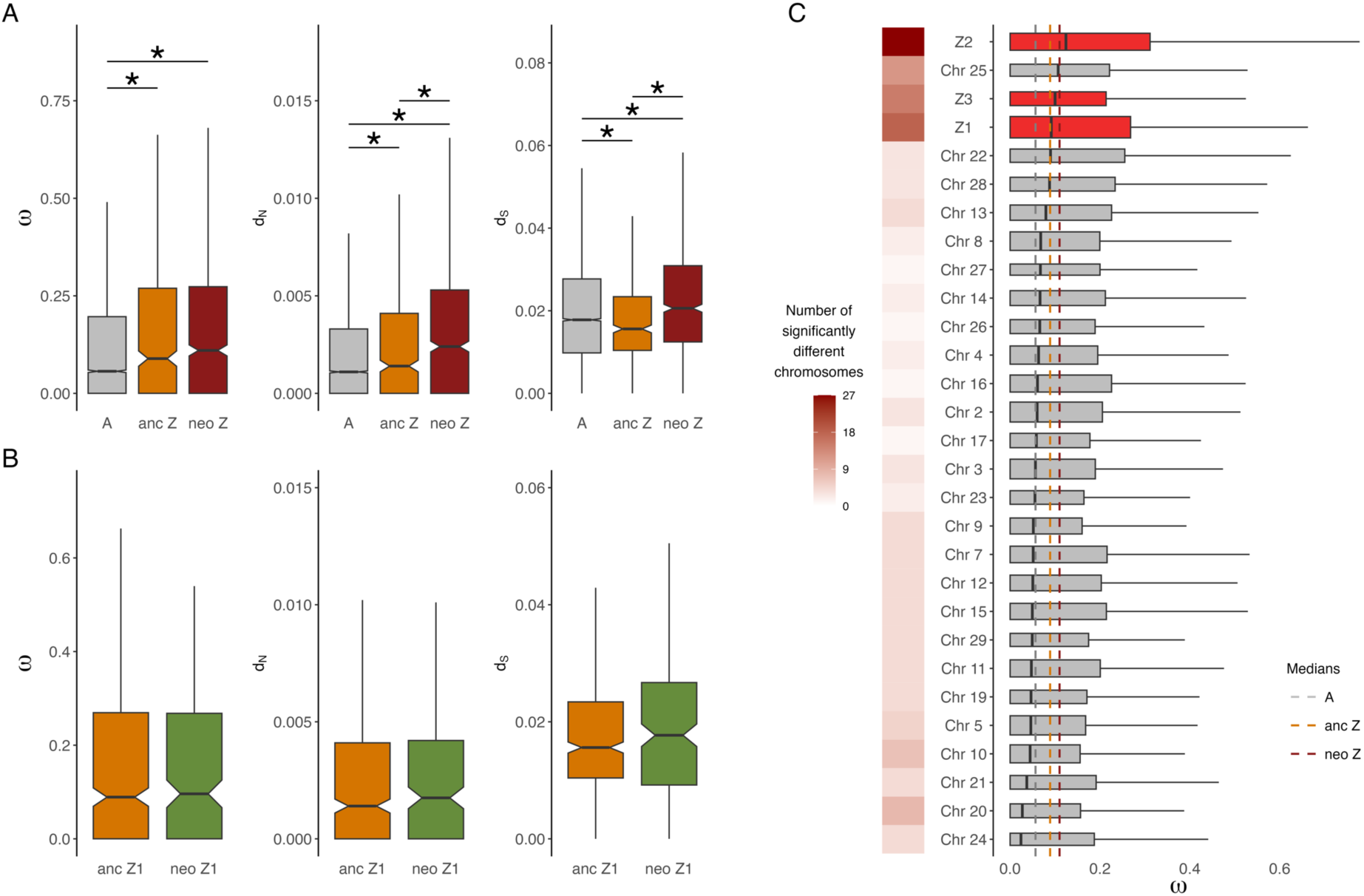
**A,** Distributions of χο, d_N_ and d_S_ for genes located on autosomes (A), the ancestral Z chromosome (anc Z) and the neo-Z chromosomes in L. sinapis. Asterisks show significant differences between categories. **B,** Distributions of χο, d_N_ and d_S_ for genes located on the ancestral and neo part of Z1 (all comparisons non-significant). **C,** Result of Dunn’s test (pair-wise comparison of all chromosomes) summarized as a heatmap showing the number of significant tests per chromosome (p-values are shown in **Supplementary table 4**). Boxplots show χο distributions for each chromosome, ordered by the median level, and dashed lines show median for each chromosome class (autosomes = A, Ancestral Z chromosome = anc Z, neo-Z chromosomes = neo Z).

**Table 1.** Median μ, *d_N_* and *d_S_* for genes located on autosomes, the ancestral Z chromosome and the neo-Z chromosomes.

| Chromosome | Number of genes | $\omega$ | $d_N$ | $d_S$ |
| --- | --- | --- | --- | --- |
| Autosomes | 9,352 | 0.0566<br>(0.0525-0.0608) | 0.0011<br>(0.0010-0.0012) | 0.0178<br>(0.0175-0.0181) |
| Ancestral Z | 477 | 0.0891<br>(0.0681-0.1111) | 0.0014<br>(0.0011-0.0018) | 0.0156<br>(0.0148-0.0168) |
| Neo-Z | 907 | 0.1101<br>(0.0986-0.1203) | 0.0024<br>(0.0022-0.0026) | 0.0206<br>(0.0193-0.0213) |

Notes. 95% confidence intervals (numbers in brackets) were obtained by bootstrapping.

**Table 2.** Median μ for genes with sex-biased or unbiased expression for the autosomes and the three different Z chromosomes.

| Sex-bias | Linkage | Number of genes | $\omega$ |
| --- | --- | --- | --- |
| **Female-biased** | Autosomes | 1,151 | 0.0625 (0.0490-0.0725) |
|  | Z1 | 34 | 0.2651 (0.0712-0.3891) |
|  | Z2 | 30 | 0.2421 (0.1043-0.3428) |
|  | Z3 | 41 | 0.1083 (0.0628-0.1362) |
| **Male-biased** | Autosomes | 1,153 | 0.0721 (0.0597-0.0800) |
|  | Z1 | 149 | 0.0556 (0.0001-0.0959) |
|  | Z2 | 85 | 0.1545 (0.1173-0.2360) |
|  | Z3 | 47 | 0.0845 (0.0671-0.1486) |
| **Unbiased** | Autosomes | 5,889 | 0.0518 (0.0474-0.0562) |
|  | Z1 | 405 | 0.0966 (0.0689-0.1247) |
|  | Z2 | 255 | 0.1010 (0.0788-0.1321) |
|  | Z3 | 161 | 0.1034 (0.0821-0.1210) |

Notes. 95% confidence intervals (numbers in brackets) were obtained by bootstrapping.

**Table 3.** Estimates of selection (median DoS and Tajima’s D) for sex-biased genes on the autosomes and the Z chromosomes.

| Statistic | Chromosome | Female-biased | Male-biased | Unbiased |
| --- | --- | --- | --- | --- |
| **DoS** | Autosomes | -0.073<br>(-0.099, -0.055) | -0.131<br>(-0.147, -0.115) | -0.100<br>(-0.111, -0.092) |
|  | All Z | -0.044<br>(-0.119, 0.000) | -0.111<br>(-0.147, -0.065) | -0.044<br>(-0.070, -0.011) |
|  | Z1 | 0.000<br>(-0.110, 0.145) | -0.114<br>(-0.177, -0.031) | -0.012<br>(-0.070, 0.000) |
|  | Z2 | 0.000<br>(-0.044, 0.086) | -0.120<br>(-0.172, -0.021) | -0.041<br>(-0.083, 0.000) |
|  | Z3 | -0.133<br>(-0.338, -0.077) | -0.087<br>(-0.150, 0.000) | -0.115<br>(-0.143, -0.024) |
| **Tajima's D** | Autosomes | -0.323<br>(-0.393, -0.257) | -0.341<br>(-0.407, -0.285) | -0.330<br>(-0.362, -0.300) |
|  | All Z | -0.749<br>(-0.967, -0.528) | -0.415<br>(-0.554, -0.287) | -0.637<br>(-0.748, -0.546) |
|  | Z1 | -0.792<br>(-1.269, -0.591) | -0.563<br>(-0.775, -0.378) | -0.722<br>(-0.813, -0.592) |
|  | Z2 | -0.734<br>(-1.141, -0.337) | -0.188<br>(-0.485, 0.070) | -0.503<br>(-0.667, -0.357) |
|  | Z3 | -0.528<br>(-0.969, -0.325) | -0.336<br>(-0.566, -0.030) | -0.656<br>(-0.932, -0.512) |

Notes. 95% confidence intervals were obtained by bootstrapping. P-values for all multiple and pair-wise comparisons between categories are presented in **Supplementary tables 9 & 10**.

Next, we analysed μ for each chromosome separately and found an overall significant difference (Kruskal-Wallis test, *χ^2^* = 139.18, p-value < 2.20*10^-16^) (**Figure 2C**). A pair-wise comparison showed that genes on Z2 had significantly higher χο compared to all other chromosomes except autosome 25. Genes on Z1 and Z3 had significantly higher χο than 16 and 13 other chromosomes respectively (**Figure 2C**, **Supplementary table 4**).

Both gene expression levels and recombination rates are expected to differ between the Z chromosomes and the autosomes and have been shown to corelate with estimates of μ (Nguyen et al., 2015; Rousselle et al., 2019). To assess the impact of these factors on the faster-Z effect, we used a linear model with μ as response variable and chromosome type, expression level and GC% at four-fold degenerate sites (as a proxy for recombination rate) as predictors. The results showed that μ had a significant positive association with both ancestral- and neo Z-linkage (p-value < 3.75*10^-3^), and a negative association with GC content (p-value < 2.2*10^-16^, **Supplementary table 5**) indicating an association between recombination rate and increased purifying selection. Expression level, however, did not have any significant effect on μ (p-value = 0.747), which can be a consequence of an underrepresentation of highly conserved genes (i.e. genes without non-synonymous substitutions that have been filtered out) which are expected to be highly/broadly expressed. Alternatively, this pattern could emerge if Z-linked genes are upregulated by dosage compensation. Regardless, these results show that there is still a strong support for a faster-Z effect after controlling for differences in expression level and GC-content at four-fold degenerate sites.

### Increased rates of divergence for female biased genes on older Z-linked regions

If the observed faster-Z effect has been caused by selection on recessive adaptive mutations in hemizygous females, we expect genes with female specific functions to have accumulated more non-synonymous substitutions. To test this prediction, we first contrasted different categories of sex-biased genes separately for each Z chromosome. This showed that female-biased genes on Z1 had a higher μ than both male-biased and unbiased genes (Wilcoxon tests, p < 0.032; **Supplementary table 6**, **Figure 3A**). Conversely, for Z2 and Z3 there were no differences in μ between any expression category (Kruskal-Wallis tests, p > 0.091; **Supplementary table 6**, **Figure 3A**). However, visual inspection of the distributions of μ for sex-biased genes on the older and the more recently acquired regions of Z1 and Z2 separately showed that female-biased genes on the older part of Z2 (corresponding to *B. mori* autosomes 24+7) appear to have evolved at a similar rate as female-biased genes on Z1 (**Figure 3A**). However, the number of female-biased genes in each region were too few for statistical comparison (n = 22, 12, 17 and 8). On the autosomes, male-biased genes had significantly higher μ compared to unbiased genes (Wilcoxon test, *W* = 3.58 × 10^6^, p = 0.007), but no other comparison was statistically significant (**Supplementary table 6**, **Figure 3A**).

**Figure 3.**
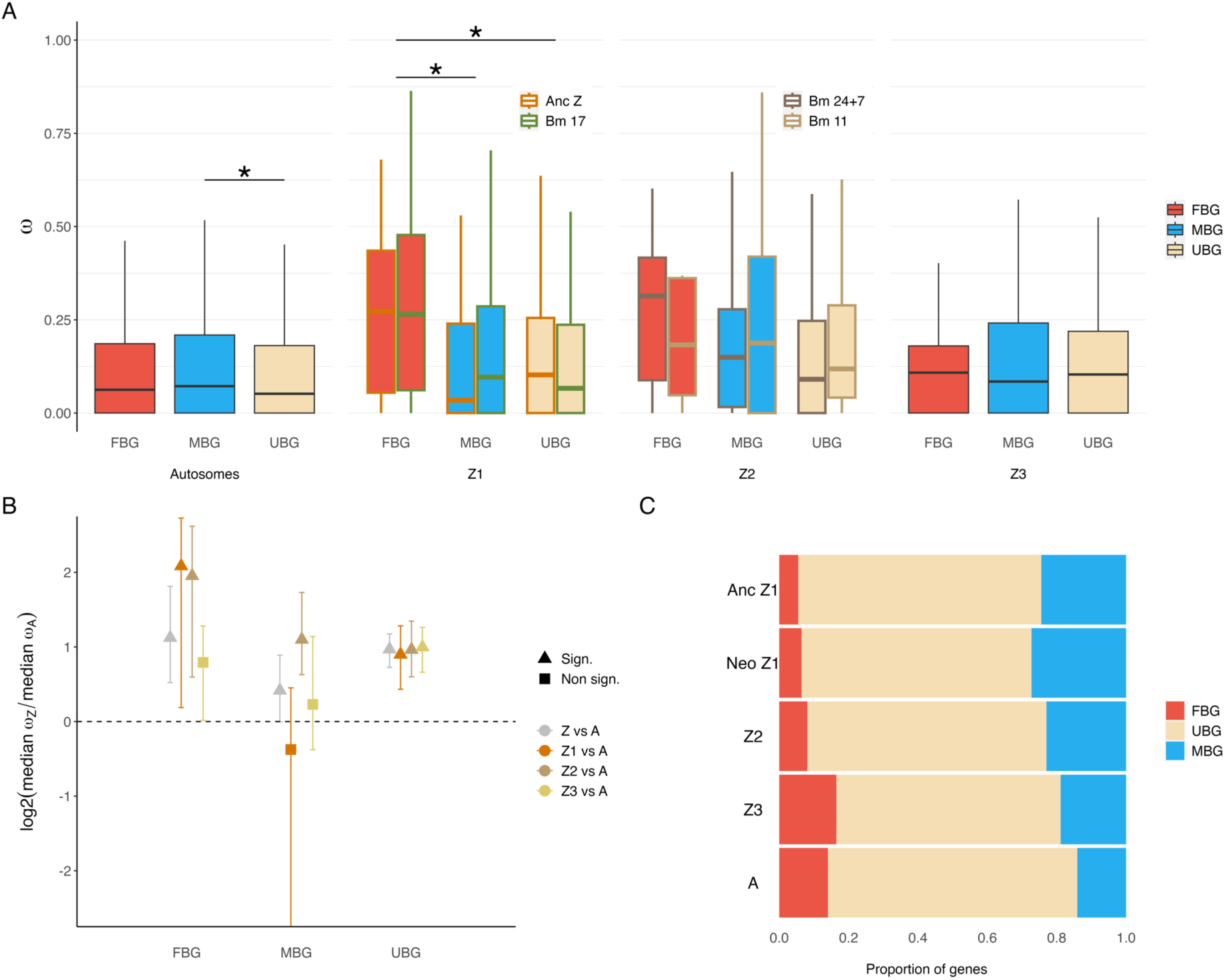
**A.** Comparison of ω between genes with sex-biased or unbiased expression for the autosomes and each Z chromosome, respectively. The ancestral (Anc Z) and neo (Bm 17) part of Z1, and the older (Bm 24+7) and the more recent (Bm 11) part of Z2 are plotted separately. **B,** Z / autosome (A) ratios of median ω for the three different Z chromosomes (Z1, Z2, Z3; log2 scaled). Triangles denote a significant difference in ω between Z-linked and autosomal genes for each expression category. Confidence intervals were obtained by bootstrapping. **C,** Proportions of sex-biased genes on the autosomes and the three different Z chromosomes. Z1 has been divided into the ancestral part (Anc Z1) and the neo part (Neo Z1). In all panels FBG = female-biased genes, MBG = male-biased genes and UBG = unbiased genes.

Next, we compared μ for Z-linked and autosomal genes with a specific sex-bias (e.g. female-biased Z-linked genes versus female-biased autosomal genes) for each Z chromosome separately (**Figure 3B**). This showed that female-biased genes had higher μ for both Z1 and Z2 (Wilcoxon tests, p < 0.002; **Supplementary table 7**), but not for Z3 (Wilcoxon test, p = 0.810; **Supplementary table 7**), as compared to autosomal female-biased genes. Male-biased genes had higher μ only on Z2 (Wilcoxon test, p = 0.001, **Supplementary table 7**), while unbiased genes had a higher μ on all Z chromosomes compared to unbiased autosomal genes (Wilcoxon tests, p < 0.0018; **Supplementary table 7**). In addition, we found that genes with sex-biased expression were non-randomly distributed across chromosome classes (Chi-squared test, *χ^2^* = 26.848, p-value = 2.13*10^-5^) (**Figure 3C**), with a higher proportion of male-biased and a lower proportion of female-biased genes on Z1, compared to the more recently derived Z2 and Z3. This clearly shows a temporal change in the proportions of sex-biased genes during sex chromosome evolution.

### Evidence for adaptive evolution of Z-linked genes

To analyse the underpinnings of the observed difference in evolutionary rates between autosomes and the Z chromosomes, we estimated the relative amount of adaptive evolution using the Direction of Selection (DoS) statistic. This revealed that Z1 and Z2 had a significantly higher DoS (median Z1 = −0.0456, Z2 = −0.0238) compared to the autosomes (median = −0.1039), but also compared to Z3 (median Z3 = −0.1186; Wilcoxon tests, p < 1.62 × 10^-4^; **Supplementary table 8**). There was, however, no difference in DoS between Z1 and Z2 or between Z3 and the autosomes (Wilcoxon tests, p > 0.846; **Supplementary table 8**, **Figure 4A**). As a complementary approach to detect differences in the efficacy of selection, we estimated Tajima’s D for all genes using population resequencing data for *L. sinapis*. For this estimate, we found a stronger signal of selection (reduced Tajima’s D) for all Z chromosomes compared to the autosomes (median Z1 = −0.6564, Z2 = −0.4535, Z3 = −0.5656, A = −0.3270; Wilcoxon tests, p < 0.016, **Supplementary table 8**), and in addition a significant reduction for Z1 compared to Z2 (Wilcoxon test, p = 0.008, **Supplementary table 8**, **Figure 4B**). We proceeded by estimating potential differences in the intensity of selection for sex-biased genes. This showed that male-biased genes had significantly lower DoS than unbiased genes when all Z-linked genes were analysed together (Wilcoxon test, p = 0.006, **Supplementary table 9**). A similar pattern was observed for Z1 and Z2 separately, but was only statistically significant for Z1 (Wilcoxon test, p = 0.018, **Supplementary table 9**, **Figure 4C**). Furthermore, DoS for male-biased genes was significantly reduced compared to both female- and unbiased genes on the autosomes (Wilcoxon tests, p < 0.020, **Supplementary table 9**), suggesting that male-biased genes evolve under more constraint in general. In contrast, DoS for female-biased genes was not significantly different from unbiased genes in any comparison (Wilcoxon test, p > 0.891; **Supplementary table 9**). For Tajima’s D, we only detected significant differences when comparing sex-biased categories for all Z-linked genes combined, with lower levels for female- and unbiased genes compared to male biased genes, again suggesting more constraint on male-biased Z-linked genes (Wilcoxon tests, p < 2.90*10^-2^, **Supplementary table 10**, **Figure 4C**). The effective population size (*N_e_*) is expected to be lower for the Z chromosomes, which could reduce the efficacy of selection on the Z chromosomes compared to the autosomes. We estimated *N_e_* based on genetic diversity at 4-fold degenerate sites (τχ_4_), and quantified the relative *N_e_* of the different Z chromosomes as τχ_4_^Z^ / τχ_4_^A^. This showed that Z1 and Z3 had low relative *N_e_* (0.51 and 0.58, respectively), while Z2 had a relative *N_e_* of 0.70, which is close to the expected level (0.75) at equal sex ratios.

**Figure 4.**
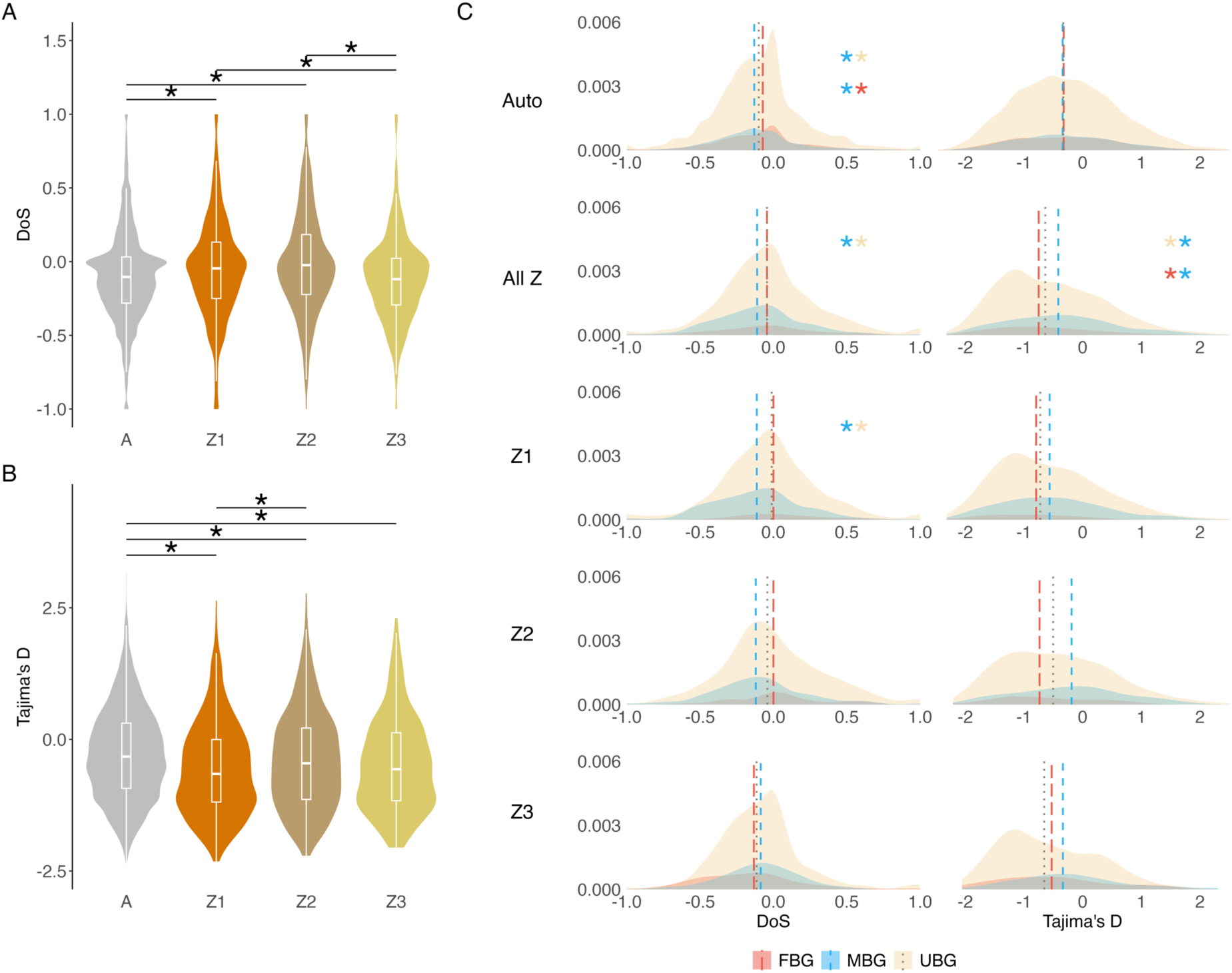
Estimates of the direction of selection (DoS) and Tajima’s D. Comparison of the distribution of DoS (**A**) and Tajima’s D (**B**) estimates for genes on the autosomes (A) and the three Z chromosomes. **C,** Distributions and comparisons of DoS and Tajima’s D estimates for female-biased (FBG), male-biased (MBG) and unbiased genes (UBG) located on autosomes (Auto), Z-chromosomes in general (All Z) and the three different Z-chromosomes, respectively (Z1, Z2, Z3). Asterisks show pair-wise comparisons that were statistically significantly different after Bonferroni correction.

## Discussion

The faster-Z/X hypothesis has been thoroughly investigated theoretically (Charlesworth et al., 1987; Mank, Vicoso, et al., 2010; Vicoso & Charlesworth, 2009) and has strong empirical support (Ávila et al., 2014; Kousathanas et al., 2014; Sackton et al., 2014; Veeramah et al., 2014). The classical explanation for this effect is that the hemizygosity of sex linked genes leads to more efficient selection, and thereby increased divergence, of sex chromosomes compared to autosomes (Charlesworth et al., 1987). While there is evidence that the efficacy of selection is higher on the sex chromosomes (Kousathanas et al., 2014; Sackton et al., 2014; Veeramah et al., 2014), the patterns vary between lineages and appear to be more ambiguous in female heterogametic systems (Mank, Nam, et al., 2010; Rousselle et al., 2016; Vicoso et al., 2013; Wright et al., 2015). Due to methodological detection biases, old and well-differentiated sex chromosomes have traditionally received more attention, while the evolutionary dynamics of newly formed sex chromosomes has been less well explored, especially in female heterogametic systems.

Although the acquisition of neo-Z chromosomes appears to be relatively common in Lepidoptera (Wright et al., 2023), little is known about their potential roles in divergence processes. The *Leptidea* clade has an unusual sex chromosome configuration with multiple neo-sex chromosomes (Šíchová et al., 2015), which makes the system particularly interesting for investigating the patterns of neo-sex chromosome evolution. An accurate dating of the neo-Z chromosomes in *Leptidea* has previously been unavailable. Here we found support for a step-wise acquisition and could narrow down the timing of two of these events through comparative inference. We found that both the neo part of Z1 and ∼ 60% of Z2 (containing homology to parts of *B. mori* autosomes 24 and 7) occur in all species and must therefore have been Z-linked > 13 MYA (Wiemers et al., 2020). The remaining ∼ 40% of Z2 (containing homology to parts of *B. mori* autosome 11) was acquired between ∼ 3-13 MYA, after the split between *L. duponcheli* and the other *Leptidea* species (Wiemers et al., 2020). The most recently acquired Z chromosome, Z3, is apparently unique to the cryptic wood white clade (*L. sinapis*, *L. juvernica* and *L. reali*), suggesting formation ∼ 3-4 MYA. Our analysis showed that a majority of Z3 genes have non-degraded neo-W gametologs, but that differentiation is substantial, resulting in an increased SNP density. These results are largely in agreement with previous qPCR results which showed that 9 out of 12 analysed genes on Z3 had equal or higher gene dose in females (Yoshido et al., 2020). This temporal variation in acquisition and levels of differentiation provides a unique opportunity to study the dynamics of butterfly sex chromosome evolution and divergence at both relatively early and late stages of sex chromosome differentiation.

In general agreement with the faster-Z hypothesis, we found an elevated rate of divergence of Z-linked genes in *L. sinapis*. This result was consistent after correcting for variation in nucleotide composition and gene expression which can have a substantial impact on patterns of divergence (Duret & Mouchiroud, 2000; Nguyen et al., 2015; Rousselle et al., 2019). A recent analysis of the butterfly *D. plexippus* showed that the neo-Z part fused with the ancestral Z, had a higher divergence rate than the ancestral Z, which in turn did not differ from the autosomes (Mongue et al., 2022). Our data did not show a significant difference in μ between genes on the ancestral Z and all neo Z-linked genes combined. Instead, more subtle differences were revealed by analysing the different Z chromosomes separately. We found the highest μ for genes on Z2, likely caused by both positive selection on female-biased genes and relaxed purifying selection for male-biased genes. In comparison, the youngest neo-Z chromosome, Z3, did not have significantly higher μ than Z1. The most straightforward explanation for the observed difference between the neo-Z chromosomes is that Z3 could have been recruited just before the radiation of the cryptic wood whites (*L. sinapis*, *L. reali* and *L. juvernica*) and therefore accumulated significantly fewer substitutions compared to the older sex chromosomes. Another obvious difference between the sex chromosomes is the level of degeneration of W-linked gametologs. This process must have started more recently for Z3 and is apparently still ongoing since we found evidence for a large proportion of W gametologs. Female-biased genes showed relatively low divergence on Z3 compared to Z1 and Z2. This is expected since a large proportion of the gametologs on neo-W might still be functional and can thereby mask recessive mutations from selection (Mrnjavac et al., 2023). Presence of functional gametologs has previously been shown to affect rates of divergence on the neo-X chromosome in *Drosophila miranda* (Zhou & Bachtrog, 2012). Conversely, for Z2, the differentiation process is complete and comparable to the state of the neo-Z in *D. plexippus* (Mongue et al., 2017), which can explain why both these chromosomes show a higher rate of divergence. In addition, the elevated genetic diversity of Z2 compared to the other Z chromosomes could contribute to making selection more efficient for this chromosome, but the reason for the difference in *N_e_* requires further study. Our results therefore support that neo-Z chromosomes can experience a temporary increase in divergence (Mongue et al., 2022), but also suggest that this requires substantial differentiation of gametologs. These observations motivate further characterization of genomic properties that could underlie the observed differences in patterns of divergence for the different Z chromosomes.

Female-biased genes are expected to be most affected by the faster-Z effect, but male-biased genes could also potentially evolve faster due to less constraints (Meisel & Connallon, 2013). We therefore explored the composition and divergence of sex-biased genes, which could be an important source of variation between the different Z chromosomes in *Leptidea*. This revealed an increased enrichment of male-biased genes (masculinization) on the Z chromosomes over time, a pattern that corroborates findings in *D. plexippus* (Mongue et al., 2022) and equivalent feminization of the *D. miranda* neo-X (Nozawa et al., 2016). This result is in line with the hypothesis that gene content often evolves towards a stronger sex-bias as the sex chromosome pair gets more differentiated (Ellegren & Parsch, 2007). In the case of *Leptidea*, the high proportion of female-biased genes on Z3 could perhaps be caused by expression of W gametologs, but this explanation requires further analysis. Furthermore, we found faster evolution of female-biased genes and particularly constrained evolution of male-biased genes in older Z chromosome regions, which have contributed to the relatively modest divergence rate on Z1. Surprisingly, this pattern was almost reversed in *D. plexippus*, where male-biased genes tended to evolve faster on the ancestral Z compared to the neo-Z chromosome (Mongue et al., 2022). Our analyses also showed that sex-biased genes had higher rates of divergence on Z2 than on Z3. In fact, genes with sex-biased expression located on Z3 did not evolve faster than their autosomal counterparts. Despite the complexity of these results, they clearly show that the variation in distribution and divergence of genes with sex-biased expression can to a large extent explain the observed differences between the three Z chromosomes. In contrast, unbiased Z-linked genes had an equally elevated divergence across all Z chromosomes compared to unbiased autosomal genes. Since this class of genes is comparatively large, they are important contributors to the faster-Z effect in *Leptidea*. There are several reasons why this pattern could emerge. Z-linked unbiased genes will experience the effect of hemizygosity when expressed in females and selection on these genes could in many cases be beneficial for both sexes (Meisel & Connallon, 2013). In addition, expression divergence can evolve rapidly for Z-linked sex-biased genes (Huylmans et al., 2017), and could thus lead to an underestimation of the effect of female-specific expression on sequence divergence (Pinharanda et al., 2019). For example, a subset of genes that historically have been female-biased, could over time have shifted to an unbiased expression pattern, leaving the signs of positive selection in the coding sequence. This explanation is in line with the observed de-feminization of the *Leptidea* Z chromosomes, if that process is mainly caused by evolution of gene expression and not by translocations.

Previous investigations of the faster-Z effect in different Lepidoptera lineages show mixed results. Increased Z chromosome divergence has been observed in *Bombyx mori* (Sackton et al., 2014), *Papilio machaon/xuthus* (Huylmans et al., 2017), *Danaus plexippus* and *Manduca sexta* (Mongue et al., 2022), but not in *Maniola jurtina*, *Pyronia tithonus* (Rousselle et al., 2016) or *Heliconius melpomene/erato* (Pinharanda et al., 2019), although Z-linked genes in *Heliconius* seem to have been more influenced by positive selection than autosomal genes (Pinharanda et al., 2019). There have also been contradictory results regarding which categories of sex-biased genes evolve faster in different species. In line with the faster-Z hypothesis, higher χο was predominantly caused by positive selection on female-biased genes on the ancestral Z chromosome in *B. mori* (Sackton et al., 2014) and on the neo-Z chromosome in *D. plexippus* (Mongue et al., 2022). In *M. sexta*, all gene categories contributed equally, while in *Papilio* butterflies, Z-linked genes with unbiased expression were mainly responsible for the faster-Z effect. It should be noted that the proportions of Z-linked sex-biased genes differed considerably between *P. machaon* and *P. xuthus*, with unusually few male-biased genes on the Z chromosome in *P. machaon* (Huylmans et al., 2017). Although the lepidopteran Z chromosome is generally enriched for male-biased genes, the relative proportions of sex-biased genes vary between species (Rousselle et al., 2016; Huylmans et al., 2017; Mongue et al., 2022), and so far, there is no obvious association between the amount of Z chromosome masculinization and the strength of the faster-Z effect. Nevertheless, our results in *Leptidea* suggest that the composition of sex-biased genes could influence the strength of the faster-Z effect, but the mechanisms and generality of this pattern requires further study.

To get more information about the underlying mechanisms driving differences in divergence between the sex chromosomes and the autosomes, we estimated summary statistics of selection. In general, we found that genes on the Z chromosomes had significantly higher DoS and reduced Tajima’s D compared to the autosomes, with the exception of Z3 for which DoS was not different from the autosomes. Our results therefore corroborate the view that Z-linked genes tend to evolve with more relaxed constraint than autosomal genes in Lepidoptera (Mongue et al., 2022; Pinharanda et al., 2019; Sackton et al., 2014). As expected, selection was pronounced for female-biased genes and, similar to divergence, appears to increase with Z chromosome age and level of differentiation. A high proportion of female-biased genes on Z3, which experience a reduced efficacy of selection, can therefore largely explain why this chromosome has a relatively low DoS compared to Z1 and Z2. This result indicates an effect of sheltering from selection by W gametologs. In contrast to female-biased genes, selection appears to have been less pronounced for male-biased genes on the Z chromosomes, although the low μ on Z1 suggests that a subset of male-biased genes could be under strong purifying selection. This is supported both by the comparatively low DoS estimates, indicating less efficient positive selection, and higher Tajima’s D estimates, implying less efficient purifying selection on recessive mutations. In addition, male-biased genes had elevated μ and reduced DoS on the autosomes compared to the other gene categories, potentially caused by a higher variance in male reproductive success which could contribute to the patterns observed for the Z chromosomes. In birds and snakes, increased Z-linked divergence is mainly explained by increased drift due to the small *N_e_* compared to the autosomes (Mank, Nam, et al., 2010; Vicoso et al., 2013; Wright et al., 2015). Effective population sizes are expected to be considerably larger in Lepidoptera than in vertebrates (Mackintosh et al., 2019), which could mean that the reduced *N_e_* of the Z chromosomes compared to the autosomes observed in *L. sinapis* (0.51 - 0.70) has a marginal effect, but could still have contributed to the observed difference in divergence between the Z chromosomes. It is therefore not unlikely that variation in effective population sizes could contribute to the mixed support for a faster-Z effect in different lepidopteran species.

The gradual degeneration of the W chromosomes should promote the evolution of dosage compensation (Wright et al., 2016). This could influence the strength of the faster-Z effect, for example by differences in expression levels (Charlesworth et al., 1987) or by gene inactivation, causing functional hemizygosity also in the homozygous sex (Mank, Vicoso, et al., 2010). In *D. plexippus*, the ancestral Z, which did not show a faster-Z effect, has the more common form of lepidopteran dosage compensation with down-regulation of Z-linked expression in males (Gu et al., 2019; Mongue et al., 2022). Conversely, the neo-Z part, which experienced a significant faster-Z effect, has evolved an upregulating mechanism in females (Gu et al., 2019; Mongue et al., 2022). It is thus possible that different mechanisms of dosage compensation occur even within chromosomes and this could potentially contribute to differences in rates of divergence. We did not find any association between expression level and divergence, which could be an indication that many of the rapidly evolving neo Z-linked genes are expressed at autosomal levels. The ancestral Z chromosome is downregulated in *L. sinapis* males (Höök et al., 2019), but dosage compensation on the neo-Z chromosomes require further investigation.

In conclusion, the step-wise addition of neo-Z chromosome regions in *Leptidea* has resulted in clear differences in how the Z-linked genes evolve. The evolution of more recently acquired neo-Z chromosome regions appears to be less influenced by positive selection. In contrast, both positive and negative selection seems to have been more at play for older Z-linked regions, in agreement with the theoretical predictions from the faster-Z hypothesis. Midway between these phases, however, there appears to be a period where a combination of relaxed selective constraints and increased positive selection can result in particularly fast divergence as observed for the *L. sinapis* Z2. The transition between these temporal phases of neo-sex chromosome evolution appears to be associated with changes in sex-biased gene expression, gradual degeneration of W-linked gametologs, and potentially influenced by the evolution of chromosome-specific dosage compensation. Further taxonomic sampling of species with more or less differentiated neo-sex chromosomes could reveal the generality of the patterns observed in *Leptidea*.

## Data availability

All raw sequencing reads have been deposited at the European Nucleotide Archive under accession PRJEB66418. All scripts used in the analysis are available at GitHub (https://github.com/EBC-butterfly-genomics-team/Leptidea_Faster_Z).

## Acknowledgements

This work was funded by the Swedish Research Council (VR research grant #019-04791 to N.B.) and by NBIS/SciLifeLab long-term bioinformatics support (WABI to N.B.). The authors acknowledge support from the National Genomics Infrastructure in Stockholm funded by Science for Life Laboratory, the Knut and Alice Wallenberg Foundation and the Swedish Research Council, and SNIC/Uppsala Multidisciplinary Center for Advanced Computational Science for assistance with massively parallel sequencing and access to the UPPMAX computational infrastructure. The computations were enabled by resources provided by the National Academic Infrastructure for Supercomputing in Sweden (NAISS) and the Swedish National Infrastructure for Computing (SNIC) in Uppsala, partially funded by the Swedish Research Council through grant agreements no. 2022-06725 and no. 2018-05973. R.V. was supported by Grant PID2022-139689NB-I00 funded by MCIN/AEI/ 10.13039/501100011033 and by ERDF A way of making Europe. We thank Jesper Boman for helpful discussions and Vlad Dincă for providing the samples for *L. morsei*, *L. amurensis*, *L. duponcheli* and *L. lactea*.

## Notes

### Competing Interest Statement

The authors have declared no competing interest.

