## Supplementary information for "Temporal Dynamics of Faster Neo-Z Evolution in Butterflies"

### Supplementary material for Temporal dynamics of faster neo-Z evolution in butterflies

#### Supplementary figures

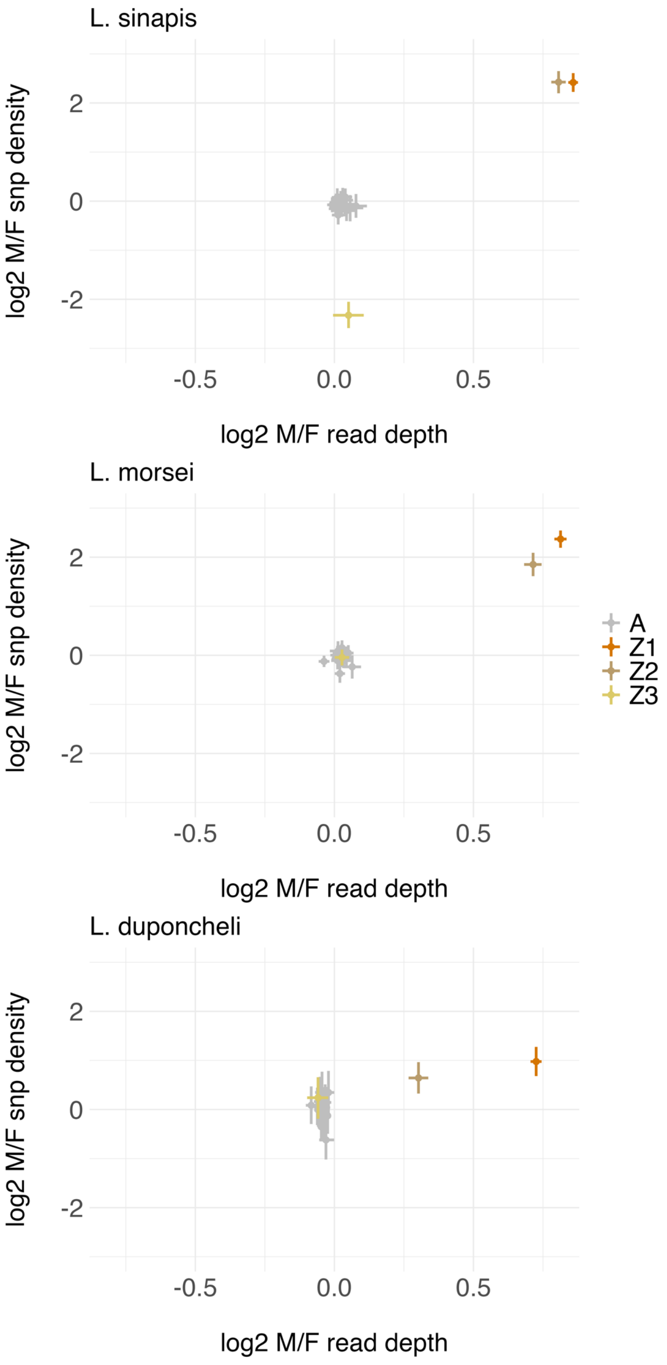


***Supplementary figure 1.*** *Mean of log2 transformed male / female ratios of read depth (gene-wise) and SNP density (100 kb windows) for each separate autosome (A) and the Z chromosomes (Z1-Z3), based on mapping against the L. sinapis reference. Lines show confidence intervals obtained by bootstrapping.*

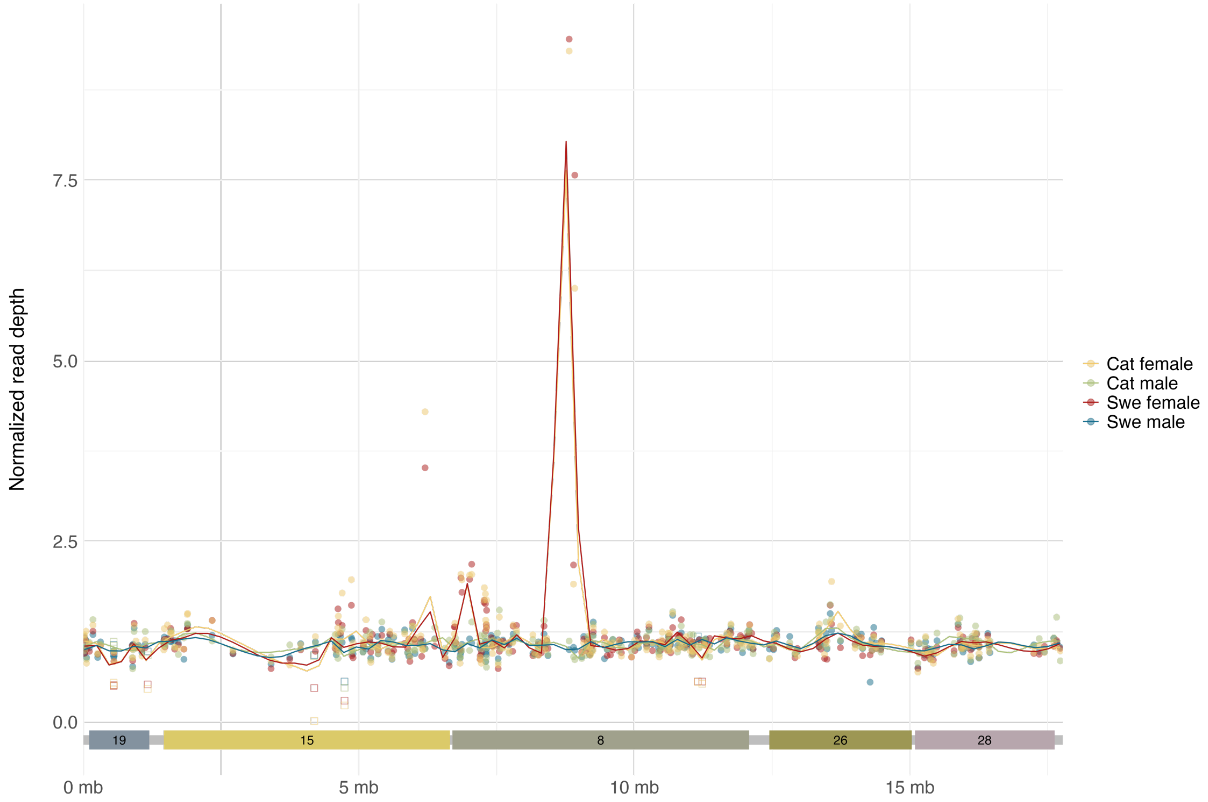


***Supplementary figure 2.*** *Normalized read depth for genes on Z3 based on mapping of two male and two female samples (one from the Swedish population and one from the Catalan population in each case). Filled dots show genes which have >80% of the L. sinapis reference male read depth across all samples and SNPs called in both female samples, indicating Z-linked genes with putative W gametologs. Empty squares show genes with reduced female read depth and SNPs not present in any of the female samples.*

A

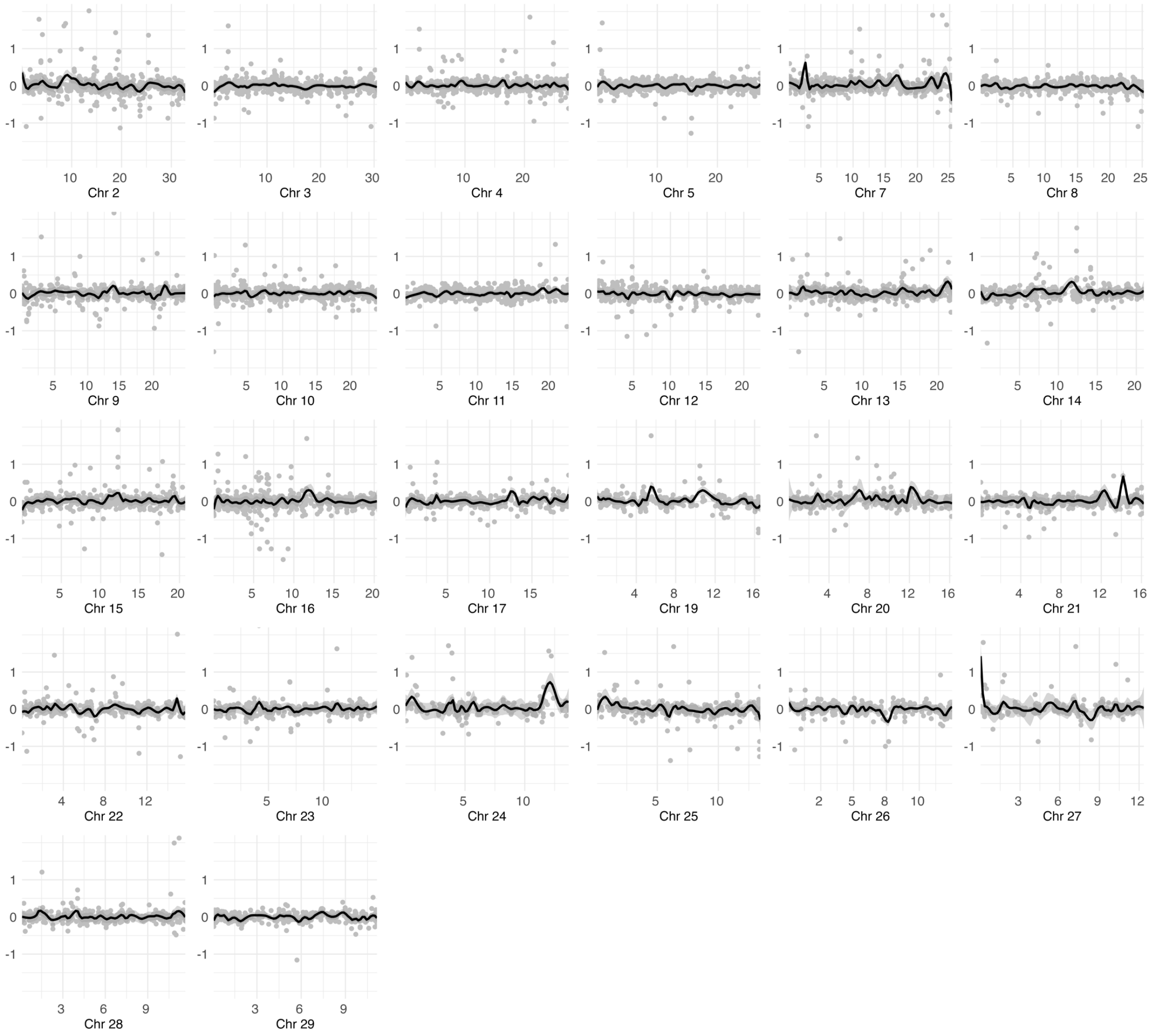
Chr 2
Chr 3
Chr 4
Chr 5
Chr 7
Chr 8
Chr 9
Chr 10
Chr 11
Chr 12
Chr 13
Chr 14
Chr 15
Chr 16
Chr 17
Chr 19
Chr 20
Chr 21
Chr 22
Chr 23
Chr 24
Chr 25
Chr 26
Chr 27
Chr 28
Chr 29

B

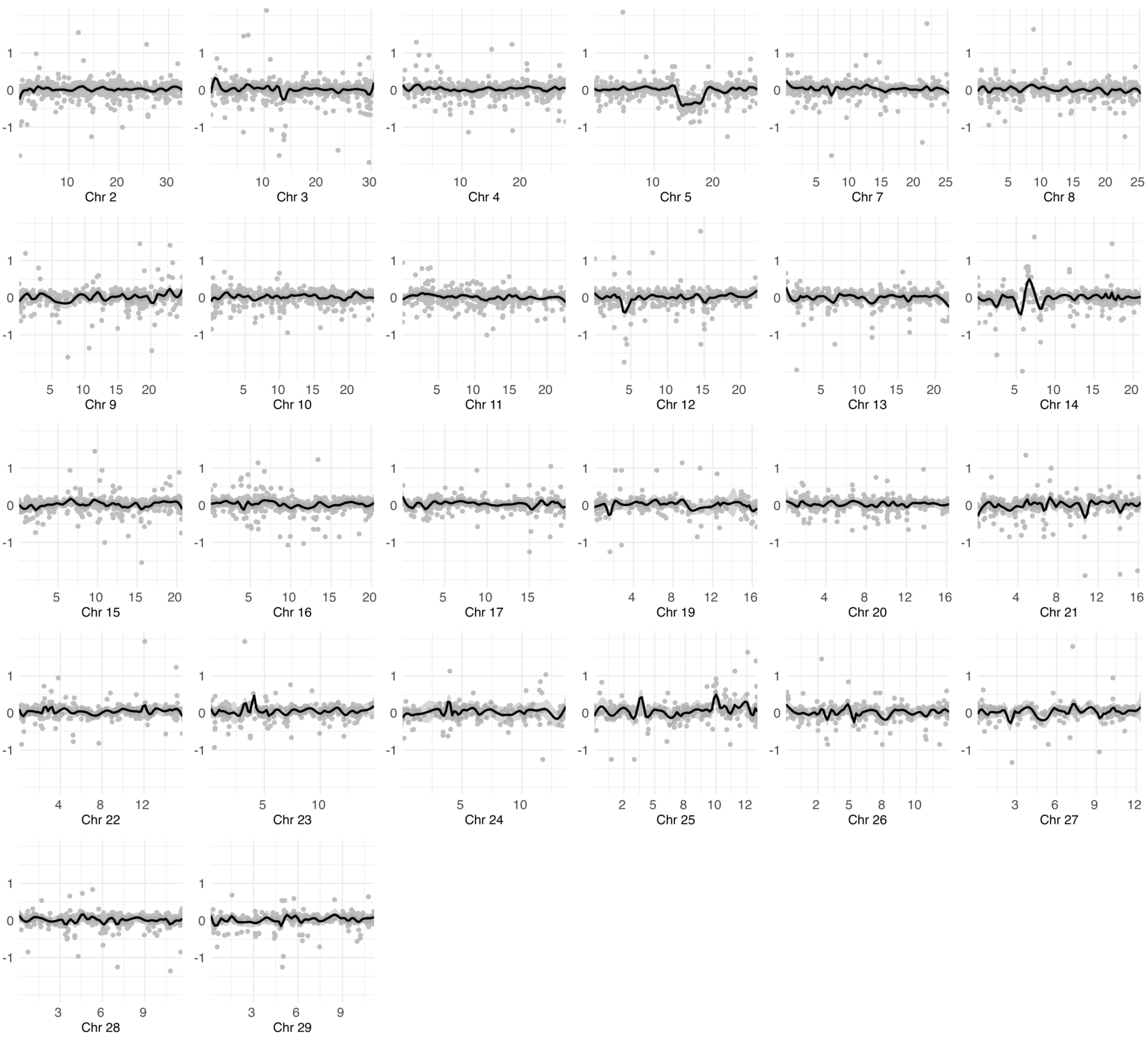
Chr 2
Chr 3
Chr 4
Chr 5
Chr 7
Chr 8
Chr 9
Chr 10
Chr 11
Chr 12
Chr 13
Chr 14
Chr 15
Chr 16
Chr 17
Chr 19
Chr 20
Chr 21
Chr 22
Chr 23
Chr 24
Chr 25
Chr 26
Chr 27
Chr 28
Chr 29

C

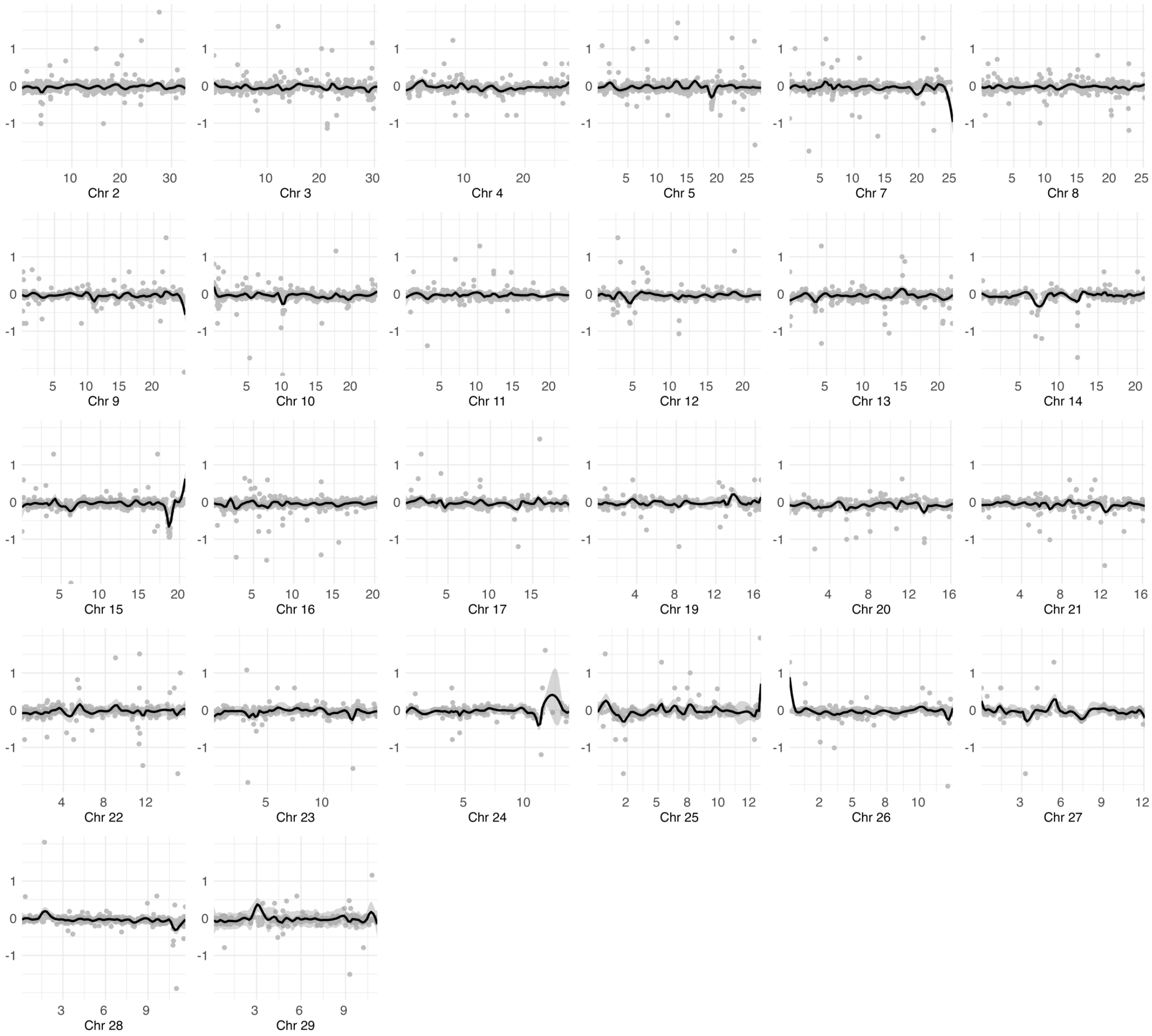


***Supplementary figure 3.*** *Log2 transformed male / female gene-wise read depth ratios across autosomes for* ***A,*** *L. sinapis;* ***B,*** *L. morsei and* ***C,*** *L. duponcheli. The X-axis scale is in Mb.*

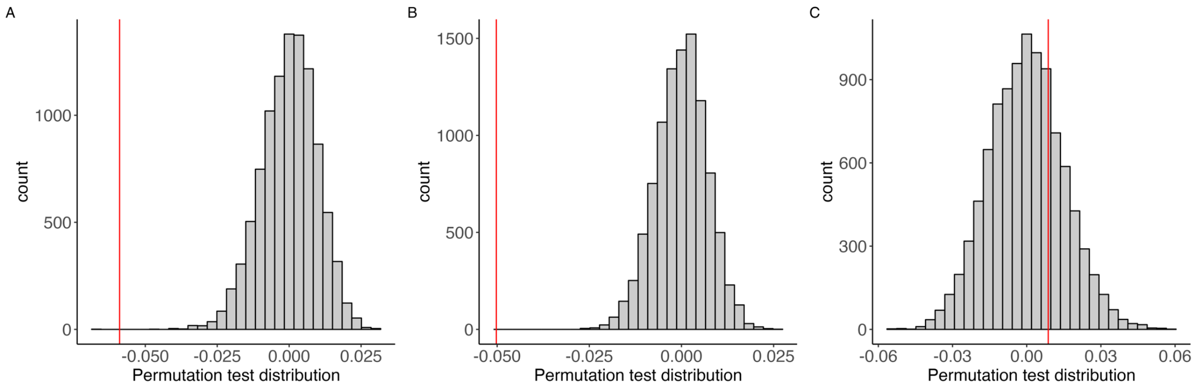


***Supplementary figure 4.*** *Permutation test results for comparing point estimates of* ω *between A, autosomes and ancestral Z; B, autosomes and neo Z; and C, ancestral and neo Z. The histograms show distributions of differences in point estimates of* ω *between randomly reshuffled groups and the red line shows the observed difference for each comparison. The tests in A and B were statistically significant after Bonferroni correction* ($p < 6*10^{-4}$) *while C was not* ($p = 1$)*.*

#### Supplementary tables

**Supplementary table 1**. Mapping statistics of samples included in the study.

| Species | Sex | Sample id | % reads mapped | Mean read depth |
|---|---|---|---|---|
| *L. sinapis* | Male | P14502_103 | 98.55 | 100.17 |
| *L. sinapis* | Female | P14502_104 | 97.93 | 77.84 |
| *L. sinapis* (Catalan) | Male | P14502_105 | 97.86 | 77.41 |
| *L. sinapis* (Catalan) | Female | P14502_106 | 97.79 | 63.95 |
| *L. amurensis* | Male | P14458_102 | 96.94 | 63.67 |
| *L. duponcheli* | Female | P14458_103 | 88.40 | 30.30 |
| *L. duponcheli* | Male | P14458_104 | 87.63 | 33.58 |
| *L. lactea* | Male | P14458_106 | 97.14 | 64.35 |
| *L. morsei* | Female | P14458_107 | 97.54 | 56.42 |
| *L. morsei* | Male | P14458_108 | 96.85 | 63.87 |

**Supplementary table 2.** Test statistics and p-values for multiple comparisons of divergence ($d_N$, $d_S$ and $\omega$) between autosomes (A) and Z chromosome categories (anc Z and neo Z) with Kruskal-Wallis rank sum test ($\chi^2$) and pair-wise comparisons with Wilcoxon rank sum test ($W$). Bonferroni corrected p-values are presented for all multiple pairwise comparisons. Significant values are highlighted in bold.

| | $d_N$ | | $d_S$ | | $\omega$ | |
|---|---|---|---|---|---|---|
| Comparison | Test statistic | P-value | Test statistic | P-value | Test statistic | P-value |
| A vs anc Z vs neo Z | $\chi^2$ = 114.35 | **< 2.20*10$^{-16}$** | $\chi^2$ = 44.069 | **2.70*10$^{-10}$** | $\chi^2$ = 84.408 | **< 2.20*10$^{-16}$** |
| A vs anc Z | $W$ = 2.02*10$^{6}$ | **4.84*10$^{-3}$** | $W$ = 2.36*10$^{6}$ | **0.020** | $W$ = 1.94*10$^{6}$ | **2.14*10$^{-5}$** |
| A vs neo Z | $W$ = 3.42*10$^{6}$ | **< 2.20*10$^{-16}$** | $W$ = 3.78*10$^{6}$ | **1.76*10$^{-8}$** | $W$ = 3.59*10$^{6}$ | **2.93*10$^{-16}$** |
| anc Z vs neo Z | $W$ = 1.90*10$^{5}$ | **6.33*10$^{-4}$** | $W$ = 1.72*10$^{5}$ | **2.03*10$^{-9}$** | $W$ = 2.07*10$^{5}$ | 0.711 |
| Z1 anc vs Z1 neo | $W$ = 4.80*10$^{4}$ | 0.821 | $W$ = 4.48*10$^{4}$ | 0.113 | $W$ = 4.96*10$^{4}$ | 0.631 |

**Supplementary table 3.** Test statistics and p-values for multiple comparisons of ENC' (effective number of codons accounting for background nucleotide composition) between autosomes (A) and Z chromosome categories (anc Z and neo Z) with Wilcoxon rank sum test (*W*). P-values are Bonferroni corrected. Significant values are highlighted in bold.

| Comparison | Test statistic | P-value |
|---|---|---|
| A vs anc Z vs neo Z | $\chi^2 = 27.534$ | **$1.05*10^{-6}$** |
| A vs anc Z | $W = 2.70 \times 10^6$ | **$4.68*10^{-7}$** |
| A vs neo Z | $W = 7.02 \times 10^6$ | 1 |
| anc Z vs neo Z | $W = 3.75 \times 10^5$ | **$6.42*10^{-5}$** |

**Supplementary table 4.** Significant results of pair-wise comparisons of ω between chromosomes using Dunn's test corrected with the Benjamini-Hochberg false discovery rate method. Significant tests are highlighted in bold.

| Group 1 | Group 2 | Number of genes1 | Number of genes 2 | Statistic (z) | P-value | P.adj |
|---|---|---|---|---|---|---|
| Z1 | Z2 | 677 | 429 | 3.221 | 0.001 | **0.011** |
| Z1 | Z3 | 677 | 280 | -0.020 | 0.984 | 0.996 |
| Z2 | Z3 | 429 | 280 | -2.606 | 0.009 | **0.048** |
| Chr 2 | Chr 20 | 589 | 286 | -2.027 | 0.043 | 0.149 |
| Chr 2 | Chr 21 | 589 | 275 | -0.604 | 0.546 | 0.731 |
| Chr 2 | Chr 22 | 589 | 262 | 1.603 | 0.109 | 0.289 |
| Chr 2 | Chr 23 | 589 | 273 | -0.226 | 0.821 | 0.914 |
| Chr 2 | Chr 24 | 589 | 219 | -1.276 | 0.202 | 0.434 |
| Chr 2 | Chr 25 | 589 | 214 | 2.537 | 0.011 | 0.055 |
| Chr 2 | Chr 26 | 589 | 287 | 0.259 | 0.796 | 0.893 |
| Chr 2 | Chr 27 | 589 | 184 | 0.147 | 0.883 | 0.946 |
| Chr 2 | Chr 28 | 589 | 280 | 1.586 | 0.113 | 0.292 |
| Chr 2 | Chr 29 | 589 | 252 | -0.558 | 0.577 | 0.755 |
| Chr 2 | Chr 3 | 589 | 572 | -0.040 | 0.968 | 0.990 |
| Chr 2 | Chr 4 | 589 | 492 | 0.412 | 0.680 | 0.827 |
| Chr 2 | Chr 5 | 589 | 544 | -1.239 | 0.215 | 0.448 |
| Chr 2 | Chr 7 | 589 | 403 | -0.431 | 0.667 | 0.820 |
| Chr 2 | Chr 8 | 589 | 509 | 0.311 | 0.756 | 0.873 |
| Chr 2 | Chr 9 | 589 | 378 | -0.827 | 0.408 | 0.638 |
| Chr 2 | Z1 | 589 | 677 | 3.454 | 0.001 | **0.006** |
| Chr 2 | Z2 | 589 | 429 | 6.197 | $5.74*10^{-10}$ | **$3.35*10^{-8}$** |
| Chr 2 | Z3 | 589 | 280 | 2.661 | 0.008 | **0.043** |
| Chr 3 | Chr 4 | 572 | 492 | 0.447 | 0.655 | 0.815 |
| Chr 3 | Chr 5 | 572 | 544 | -1.191 | 0.234 | 0.470 |
| Chr 3 | Chr 7 | 572 | 403 | -0.392 | 0.695 | 0.837 |
| Chr 3 | Chr 8 | 572 | 509 | 0.347 | 0.728 | 0.861 |
| Chr 3 | Chr 9 | 572 | 378 | -0.787 | 0.431 | 0.666 |
| Chr 3 | Z1 | 572 | 677 | 3.468 | 0.001 | **0.005** |
| Chr 3 | Z2 | 572 | 429 | 6.196 | $5.80*10^{-10}$ | **$3.35*10^{-8}$** |
| Chr 3 | Z3 | 572 | 280 | 2.681 | 0.007 | **0.041** |
| Chr 4 | Chr 5 | 492 | 544 | -1.588 | 0.112 | 0.292 |
| Chr 4 | Chr 7 | 492 | 403 | -0.789 | 0.430 | 0.666 |
| Chr 4 | Chr 8 | 492 | 509 | -0.100 | 0.920 | 0.965 |
| Chr 4 | Chr 9 | 492 | 378 | -1.165 | 0.244 | 0.479 |
| Chr 4 | Z1 | 492 | 677 | 2.860 | 0.004 | **0.028** |
| Chr 4 | Z2 | 492 | 429 | 5.574 | $2.49*10^{-8}$ | **$6.74*10^{-7}$** |
| Chr 4 | Z3 | 492 | 280 | 2.244 | 0.025 | 0.101 |
| Chr 5 | Chr 7 | 544 | 403 | 0.697 | 0.486 | 0.700 |
| Chr 5 | Chr 8 | 544 | 509 | 1.500 | 0.134 | 0.317 |
| Chr 5 | Chr 9 | 544 | 378 | 0.286 | 0.775 | 0.879 |

| | | | | | | |
|---|---|---|---|---|---|---|
| Chr 5 | Z1 | 544 | 677 | 4.659 | $3.17*10^{-6}$ | **$5.60*10^{-5}$** |
| Chr 5 | Z2 | 544 | 429 | 7.233 | $4.73*10^{-13}$ | **$9.60*10^{-11}$** |
| Chr 5 | Z3 | 544 | 280 | 3.628 | $2.86*10^{-4}$ | **0.003** |
| Chr 7 | Chr 8 | 403 | 509 | 0.700 | 0.484 | 0.700 |
| Chr 7 | Chr 9 | 403 | 378 | -0.373 | 0.710 | 0.850 |
| Chr 7 | Z1 | 403 | 677 | 3.536 | $4.06*10^{-4}$ | **0.004** |
| Chr 7 | Z2 | 403 | 429 | 6.072 | $1.26*10^{-9}$ | **$5.70*10^{-8}$** |
| Chr 7 | Z3 | 403 | 280 | 2.841 | 0.004 | **0.029** |
| Chr 8 | Chr 9 | 509 | 378 | -1.080 | 0.280 | 0.512 |
| Chr 8 | Z1 | 509 | 677 | 2.997 | 0.003 | **0.019** |
| Chr 8 | Z2 | 509 | 429 | 5.715 | $1.10*10^{-8}$ | **$3.43*10^{-7}$** |
| Chr 8 | Z3 | 509 | 280 | 2.343 | 0.019 | 0.081 |
| Chr 9 | Z1 | 378 | 677 | 3.880 | $1.04*10^{-4}$ | **0.001** |
| Chr 9 | Z2 | 378 | 429 | 6.349 | $2.17*10^{-10}$ | **$1.76*10^{-8}$** |
| Chr 9 | Z3 | 378 | 280 | 3.142 | 0.002 | **0.013** |
| Chr 10 | Chr 11 | 480 | 430 | 0.831 | 0.406 | 0.638 |
| Chr 10 | Chr 12 | 480 | 362 | 1.012 | 0.311 | 0.545 |
| Chr 10 | Chr 13 | 480 | 344 | 2.904 | 0.004 | **0.025** |
| Chr 10 | Chr 14 | 480 | 327 | 2.352 | 0.019 | 0.080 |
| Chr 10 | Chr 15 | 480 | 386 | 1.134 | 0.257 | 0.492 |
| Chr 10 | Chr 16 | 480 | 429 | 2.135 | 0.033 | 0.124 |
| Chr 10 | Chr 17 | 480 | 306 | 1.626 | 0.104 | 0.282 |
| Chr 10 | Chr 19 | 480 | 264 | 0.129 | 0.898 | 0.954 |
| Chr 10 | Chr 2 | 480 | 589 | 1.619 | 0.105 | 0.284 |
| Chr 10 | Chr 20 | 480 | 286 | -0.623 | 0.533 | 0.729 |
| Chr 10 | Chr 21 | 480 | 275 | 0.733 | 0.463 | 0.686 |
| Chr 10 | Chr 22 | 480 | 262 | 2.845 | 0.004 | **0.029** |
| Chr 10 | Chr 23 | 480 | 273 | 1.095 | 0.273 | 0.509 |
| Chr 10 | Chr 24 | 480 | 219 | -0.017 | 0.986 | 0.996 |
| Chr 10 | Chr 25 | 480 | 214 | 3.675 | $2.38*10^{-4}$ | **0.003** |
| Chr 10 | Chr 26 | 480 | 287 | 1.584 | 0.113 | 0.292 |
| Chr 10 | Chr 27 | 480 | 184 | 1.291 | 0.197 | 0.427 |
| Chr 10 | Chr 28 | 480 | 280 | 2.855 | 0.004 | **0.028** |
| Chr 10 | Chr 29 | 480 | 252 | 0.739 | 0.460 | 0.684 |
| Chr 10 | Chr 3 | 480 | 572 | 1.570 | 0.116 | 0.297 |
| Chr 10 | Chr 4 | 480 | 492 | 1.944 | 0.052 | 0.173 |
| Chr 10 | Chr 5 | 480 | 544 | 0.413 | 0.679 | 0.827 |
| Chr 10 | Chr 7 | 480 | 403 | 1.061 | 0.289 | 0.521 |
| Chr 10 | Chr 8 | 480 | 509 | 1.860 | 0.063 | 0.196 |
| Chr 10 | Chr 9 | 480 | 378 | 0.655 | 0.513 | 0.720 |
| Chr 10 | Z1 | 480 | 677 | 4.930 | $8.22*10^{-7}$ | **$1.67*10^{-5}$** |
| Chr 10 | Z2 | 480 | 429 | 7.419 | $1.18*10^{-13}$ | **$4.80*10^{-11}$** |
| Chr 10 | Z3 | 480 | 280 | 3.893 | $9.91*10^{-5}$ | **0.001** |
| Chr 11 | Chr 12 | 430 | 362 | 0.214 | 0.830 | 0.919 |
| Chr 11 | Chr 13 | 430 | 344 | 2.073 | 0.038 | 0.138 |
| Chr 11 | Chr 14 | 430 | 327 | 1.546 | 0.122 | 0.302 |
| Chr 11 | Chr 15 | 430 | 386 | 0.319 | 0.750 | 0.872 |
| Chr 11 | Chr 16 | 430 | 429 | 1.270 | 0.204 | 0.434 |
| Chr 11 | Chr 17 | 430 | 306 | 0.852 | 0.394 | 0.632 |
| Chr 11 | Chr 19 | 430 | 264 | -0.580 | 0.562 | 0.746 |
| Chr 11 | Chr 2 | 430 | 589 | 0.699 | 0.484 | 0.700 |
| Chr 11 | Chr 20 | 430 | 286 | -1.333 | 0.183 | 0.412 |
| Chr 11 | Chr 21 | 430 | 275 | 0.004 | 0.997 | 0.998 |
| Chr 11 | Chr 22 | 430 | 262 | 2.084 | 0.037 | 0.138 |
| Chr 11 | Chr 23 | 430 | 273 | 0.360 | 0.719 | 0.855 |
| Chr 11 | Chr 24 | 430 | 219 | -0.682 | 0.495 | 0.708 |
| Chr 11 | Chr 25 | 430 | 214 | 2.951 | 0.003 | **0.022** |
| Chr 11 | Chr 26 | 430 | 287 | 0.826 | 0.409 | 0.638 |
| Chr 11 | Chr 27 | 430 | 184 | 0.645 | 0.519 | 0.727 |
| Chr 11 | Chr 28 | 430 | 280 | 2.077 | 0.038 | 0.138 |

| | | | | | | |
|---|---|---|---|---|---|---|
| Chr 11 | Chr 29 | 430 | 252 | 0.029 | 0.977 | 0.996 |
| Chr 11 | Chr 3 | 430 | 572 | 0.658 | 0.510 | 0.720 |
| Chr 11 | Chr 4 | 430 | 492 | 1.053 | 0.292 | 0.525 |
| Chr 11 | Chr 5 | 430 | 544 | -0.454 | 0.650 | 0.815 |
| Chr 11 | Chr 7 | 430 | 403 | 0.238 | 0.812 | 0.905 |
| Chr 11 | Chr 8 | 430 | 509 | 0.965 | 0.335 | 0.569 |
| Chr 11 | Chr 9 | 430 | 378 | -0.144 | 0.885 | 0.946 |
| Chr 11 | Z1 | 430 | 677 | 3.875 | $1.06*10^{-4}$ | **0.001** |
| Chr 11 | Z2 | 430 | 429 | 6.415 | $1.41*10^{-10}$ | **$1.43*10^{-8}$** |
| Chr 11 | Z3 | 430 | 280 | 3.093 | 0.002 | **0.015** |
| Chr 12 | Chr 13 | 362 | 344 | 1.789 | 0.074 | 0.214 |
| Chr 12 | Chr 14 | 362 | 327 | 1.287 | 0.198 | 0.428 |
| Chr 12 | Chr 15 | 362 | 386 | 0.097 | 0.923 | 0.966 |
| Chr 12 | Chr 16 | 362 | 429 | 1.000 | 0.317 | 0.548 |
| Chr 12 | Chr 17 | 362 | 306 | 0.624 | 0.533 | 0.729 |
| Chr 12 | Chr 19 | 362 | 264 | -0.749 | 0.454 | 0.682 |
| Chr 12 | Chr 2 | 362 | 589 | 0.435 | 0.663 | 0.820 |
| Chr 12 | Chr 20 | 362 | 286 | -1.479 | 0.139 | 0.327 |
| Chr 12 | Chr 21 | 362 | 275 | -0.188 | 0.851 | 0.934 |
| Chr 12 | Chr 22 | 362 | 262 | 1.826 | 0.068 | 0.204 |
| Chr 12 | Chr 23 | 362 | 273 | 0.157 | 0.876 | 0.946 |
| Chr 12 | Chr 24 | 362 | 219 | -0.840 | 0.401 | 0.634 |
| Chr 12 | Chr 25 | 362 | 214 | 2.685 | 0.007 | **0.041** |
| Chr 12 | Chr 26 | 362 | 287 | 0.604 | 0.546 | 0.731 |
| Chr 12 | Chr 27 | 362 | 184 | 0.458 | 0.647 | 0.815 |
| Chr 12 | Chr 28 | 362 | 280 | 1.812 | 0.070 | 0.207 |
| Chr 12 | Chr 29 | 362 | 252 | -0.158 | 0.875 | 0.946 |
| Chr 12 | Chr 3 | 362 | 572 | 0.398 | 0.691 | 0.835 |
| Chr 12 | Chr 4 | 362 | 492 | 0.783 | 0.433 | 0.666 |
| Chr 12 | Chr 5 | 362 | 544 | -0.657 | 0.511 | 0.720 |
| Chr 12 | Chr 7 | 362 | 403 | 0.017 | 0.986 | 0.996 |
| Chr 12 | Chr 8 | 362 | 509 | 0.697 | 0.486 | 0.700 |
| Chr 12 | Chr 9 | 362 | 378 | -0.346 | 0.729 | 0.861 |
| Chr 12 | Z1 | 362 | 677 | 3.436 | 0.001 | **0.006** |
| Chr 12 | Z2 | 362 | 429 | 5.919 | $3.24*10^{-9}$ | **$1.09*10^{-7}$** |
| Chr 12 | Z3 | 362 | 280 | 2.793 | 0.005 | **0.032** |
| Chr 13 | Chr 14 | 344 | 327 | -0.473 | 0.636 | 0.806 |
| Chr 13 | Chr 15 | 344 | 386 | -1.721 | 0.085 | 0.239 |
| Chr 13 | Chr 16 | 344 | 429 | -0.875 | 0.382 | 0.622 |
| Chr 13 | Chr 17 | 344 | 306 | -1.098 | 0.272 | 0.509 |
| Chr 13 | Chr 19 | 344 | 264 | -2.387 | 0.017 | 0.073 |
| Chr 13 | Chr 2 | 344 | 589 | -1.556 | 0.120 | 0.302 |
| Chr 13 | Chr 20 | 344 | 286 | -3.145 | 0.002 | **0.013** |
| Chr 13 | Chr 21 | 344 | 275 | -1.851 | 0.064 | 0.198 |
| Chr 13 | Chr 22 | 344 | 262 | 0.163 | 0.870 | 0.946 |
| Chr 13 | Chr 23 | 344 | 273 | -1.507 | 0.132 | 0.316 |
| Chr 13 | Chr 24 | 344 | 219 | -2.390 | 0.017 | 0.073 |
| Chr 13 | Chr 25 | 344 | 214 | 1.113 | 0.266 | 0.502 |
| Chr 13 | Chr 26 | 344 | 287 | -1.088 | 0.277 | 0.511 |
| Chr 13 | Chr 27 | 344 | 184 | -1.020 | 0.308 | 0.543 |
| Chr 13 | Chr 28 | 344 | 280 | 0.118 | 0.906 | 0.958 |
| Chr 13 | Chr 29 | 344 | 252 | -1.781 | 0.075 | 0.216 |
| Chr 13 | Chr 3 | 344 | 572 | -1.582 | 0.114 | 0.292 |
| Chr 13 | Chr 4 | 344 | 492 | -1.145 | 0.252 | 0.488 |
| Chr 13 | Chr 5 | 344 | 544 | -2.603 | 0.009 | **0.048** |
| Chr 13 | Chr 7 | 344 | 403 | -1.818 | 0.069 | 0.206 |
| Chr 13 | Chr 8 | 344 | 509 | -1.244 | 0.214 | 0.448 |
| Chr 13 | Chr 9 | 344 | 378 | -2.149 | 0.032 | 0.121 |
| Chr 13 | Z1 | 344 | 677 | 1.344 | 0.179 | 0.408 |
| Chr 13 | Z2 | 344 | 429 | 3.976 | $7.01*10^{-5}$ | **0.001** |

| | | | | | | |
|---|---|---|---|---|---|---|
| Chr 13 | Z3 | 344 | 280 | 1.088 | 0.277 | 0.511 |
| Chr 14 | Chr 15 | 327 | 386 | -1.212 | 0.226 | 0.463 |
| Chr 14 | Chr 16 | 327 | 429 | -0.365 | 0.715 | 0.854 |
| Chr 14 | Chr 17 | 327 | 306 | -0.625 | 0.532 | 0.729 |
| Chr 14 | Chr 19 | 327 | 264 | -1.919 | 0.055 | 0.182 |
| Chr 14 | Chr 2 | 327 | 589 | -1.002 | 0.317 | 0.548 |
| Chr 14 | Chr 20 | 327 | 286 | -2.657 | 0.008 | **0.043** |
| Chr 14 | Chr 21 | 327 | 275 | -1.383 | 0.167 | 0.387 |
| Chr 14 | Chr 22 | 327 | 262 | 0.602 | 0.547 | 0.731 |
| Chr 14 | Chr 23 | 327 | 273 | -1.044 | 0.296 | 0.530 |
| Chr 14 | Chr 24 | 327 | 219 | -1.947 | 0.051 | 0.173 |
| Chr 14 | Chr 25 | 327 | 214 | 1.517 | 0.129 | 0.312 |
| Chr 14 | Chr 26 | 327 | 287 | -0.624 | 0.533 | 0.729 |
| Chr 14 | Chr 27 | 327 | 184 | -0.615 | 0.539 | 0.731 |
| Chr 14 | Chr 28 | 327 | 280 | 0.566 | 0.571 | 0.753 |
| Chr 14 | Chr 29 | 327 | 252 | -1.325 | 0.185 | 0.415 |
| Chr 14 | Chr 3 | 327 | 572 | -1.030 | 0.303 | 0.537 |
| Chr 14 | Chr 4 | 327 | 492 | -0.615 | 0.538 | 0.731 |
| Chr 14 | Chr 5 | 327 | 544 | -2.040 | 0.041 | 0.147 |
| Chr 14 | Chr 7 | 327 | 403 | -1.302 | 0.193 | 0.426 |
| Chr 14 | Chr 8 | 327 | 509 | -0.709 | 0.478 | 0.700 |
| Chr 14 | Chr 9 | 327 | 378 | -1.637 | 0.102 | 0.279 |
| Chr 14 | Z1 | 327 | 677 | 1.864 | 0.062 | 0.196 |
| Chr 14 | Z2 | 327 | 429 | 4.417 | $9.99*10^{-6}$ | **$1.69*10^{-4}$** |
| Chr 14 | Z3 | 327 | 280 | 1.524 | 0.127 | 0.310 |
| Chr 15 | Chr 16 | 386 | 429 | 0.917 | 0.359 | 0.591 |
| Chr 15 | Chr 17 | 386 | 306 | 0.541 | 0.589 | 0.761 |
| Chr 15 | Chr 19 | 386 | 264 | -0.848 | 0.397 | 0.632 |
| Chr 15 | Chr 2 | 386 | 589 | 0.336 | 0.737 | 0.862 |
| Chr 15 | Chr 20 | 386 | 286 | -1.590 | 0.112 | 0.292 |
| Chr 15 | Chr 21 | 386 | 275 | -0.280 | 0.780 | 0.882 |
| Chr 15 | Chr 22 | 386 | 262 | 1.762 | 0.078 | 0.223 |
| Chr 15 | Chr 23 | 386 | 273 | 0.069 | 0.945 | 0.974 |
| Chr 15 | Chr 24 | 386 | 219 | -0.933 | 0.351 | 0.586 |
| Chr 15 | Chr 25 | 386 | 214 | 2.634 | 0.008 | **0.046** |
| Chr 15 | Chr 26 | 386 | 287 | 0.521 | 0.602 | 0.771 |
| Chr 15 | Chr 27 | 386 | 184 | 0.384 | 0.701 | 0.842 |
| Chr 15 | Chr 28 | 386 | 280 | 1.747 | 0.081 | 0.229 |
| Chr 15 | Chr 29 | 386 | 252 | -0.247 | 0.805 | 0.900 |
| Chr 15 | Chr 3 | 386 | 572 | 0.299 | 0.765 | 0.875 |
| Chr 15 | Chr 4 | 386 | 492 | 0.694 | 0.488 | 0.700 |
| Chr 15 | Chr 5 | 386 | 544 | -0.776 | 0.438 | 0.668 |
| Chr 15 | Chr 7 | 386 | 403 | -0.082 | 0.935 | 0.972 |
| Chr 15 | Chr 8 | 386 | 509 | 0.605 | 0.545 | 0.731 |
| Chr 15 | Chr 9 | 386 | 378 | -0.449 | 0.653 | 0.815 |
| Chr 15 | Z1 | 386 | 677 | 3.397 | 0.001 | **0.006** |
| Chr 15 | Z2 | 386 | 429 | 5.921 | $3.20*10^{-9}$ | **$1.09*10^{-7}$** |
| Chr 15 | Z3 | 386 | 280 | 2.741 | 0.006 | **0.037** |
| Chr 16 | Chr 17 | 429 | 306 | -0.307 | 0.759 | 0.873 |
| Chr 16 | Chr 19 | 429 | 264 | -1.688 | 0.091 | 0.254 |
| Chr 16 | Chr 2 | 429 | 589 | -0.666 | 0.505 | 0.720 |
| Chr 16 | Chr 20 | 429 | 286 | -2.468 | 0.014 | 0.064 |
| Chr 16 | Chr 21 | 429 | 275 | -1.118 | 0.263 | 0.500 |
| Chr 16 | Chr 22 | 429 | 262 | 0.978 | 0.328 | 0.562 |
| Chr 16 | Chr 23 | 429 | 273 | -0.760 | 0.447 | 0.675 |
| Chr 16 | Chr 24 | 429 | 219 | -1.725 | 0.085 | 0.238 |
| Chr 16 | Chr 25 | 429 | 214 | 1.914 | 0.056 | 0.182 |
| Chr 16 | Chr 26 | 429 | 287 | -0.310 | 0.756 | 0.873 |
| Chr 16 | Chr 27 | 429 | 184 | -0.339 | 0.735 | 0.862 |
| Chr 16 | Chr 28 | 429 | 280 | 0.948 | 0.343 | 0.578 |

| | | | | | | |
|---|---|---|---|---|---|---|
| Chr 16 | Chr 29 | 429 | 252 | -1.063 | 0.288 | 0.521 |
| Chr 16 | Chr 3 | 429 | 572 | -0.699 | 0.484 | 0.700 |
| Chr 16 | Chr 4 | 429 | 492 | -0.259 | 0.795 | 0.893 |
| Chr 16 | Chr 5 | 429 | 544 | -1.796 | 0.072 | 0.212 |
| Chr 16 | Chr 7 | 429 | 403 | -1.011 | 0.312 | 0.545 |
| Chr 16 | Chr 8 | 429 | 509 | -0.358 | 0.720 | 0.855 |
| Chr 16 | Chr 9 | 429 | 378 | -1.373 | 0.170 | 0.392 |
| Chr 16 | Z1 | 429 | 677 | 2.468 | 0.014 | 0.064 |
| Chr 16 | Z2 | 429 | 429 | 5.142 | $2.72*10^{-7}$ | **$6.15*10^{-6}$** |
| Chr 16 | Z3 | 429 | 280 | 1.964 | 0.050 | 0.168 |
| Chr 17 | Chr 19 | 306 | 264 | -1.298 | 0.194 | 0.426 |
| Chr 17 | Chr 2 | 306 | 589 | -0.275 | 0.784 | 0.884 |
| Chr 17 | Chr 20 | 306 | 286 | -2.011 | 0.044 | 0.154 |
| Chr 17 | Chr 21 | 306 | 275 | -0.763 | 0.445 | 0.674 |
| Chr 17 | Chr 22 | 306 | 262 | 1.184 | 0.236 | 0.473 |
| Chr 17 | Chr 23 | 306 | 273 | -0.431 | 0.666 | 0.820 |
| Chr 17 | Chr 24 | 306 | 219 | -1.359 | 0.174 | 0.399 |
| Chr 17 | Chr 25 | 306 | 214 | 2.055 | 0.040 | 0.143 |
| Chr 17 | Chr 26 | 306 | 287 | -0.009 | 0.993 | 0.998 |
| Chr 17 | Chr 27 | 306 | 184 | -0.074 | 0.941 | 0.972 |
| Chr 17 | Chr 28 | 306 | 280 | 1.158 | 0.247 | 0.482 |
| Chr 17 | Chr 29 | 306 | 252 | -0.722 | 0.470 | 0.695 |
| Chr 17 | Chr 3 | 306 | 572 | -0.307 | 0.759 | 0.873 |
| Chr 17 | Chr 4 | 306 | 492 | 0.080 | 0.936 | 0.972 |
| Chr 17 | Chr 5 | 306 | 544 | -1.302 | 0.193 | 0.426 |
| Chr 17 | Chr 7 | 306 | 403 | -0.623 | 0.534 | 0.729 |
| Chr 17 | Chr 8 | 306 | 509 | -0.008 | 0.994 | 0.998 |
| Chr 17 | Chr 9 | 306 | 378 | -0.961 | 0.337 | 0.569 |
| Chr 17 | Z1 | 306 | 677 | 2.544 | 0.011 | 0.054 |
| Chr 17 | Z2 | 306 | 429 | 4.998 | $5.78*10^{-7}$ | **$1.24*10^{-5}$** |
| Chr 17 | Z3 | 306 | 280 | 2.102 | 0.036 | 0.134 |
| Chr 19 | Chr 2 | 264 | 589 | 1.211 | 0.226 | 0.463 |
| Chr 19 | Chr 20 | 264 | 286 | -0.660 | 0.509 | 0.720 |
| Chr 19 | Chr 21 | 264 | 275 | 0.529 | 0.596 | 0.769 |
| Chr 19 | Chr 22 | 264 | 262 | 2.393 | 0.017 | 0.073 |
| Chr 19 | Chr 23 | 264 | 273 | 0.848 | 0.397 | 0.632 |
| Chr 19 | Chr 24 | 264 | 219 | -0.123 | 0.902 | 0.956 |
| Chr 19 | Chr 25 | 264 | 214 | 3.177 | 0.001 | **0.012** |
| Chr 19 | Chr 26 | 264 | 287 | 1.270 | 0.204 | 0.434 |
| Chr 19 | Chr 27 | 264 | 184 | 1.063 | 0.288 | 0.521 |
| Chr 19 | Chr 28 | 264 | 280 | 2.388 | 0.017 | 0.073 |
| Chr 19 | Chr 29 | 264 | 252 | 0.541 | 0.588 | 0.761 |
| Chr 19 | Chr 3 | 264 | 572 | 1.174 | 0.240 | 0.476 |
| Chr 19 | Chr 4 | 264 | 492 | 1.506 | 0.132 | 0.316 |
| Chr 19 | Chr 5 | 264 | 544 | 0.214 | 0.831 | 0.919 |
| Chr 19 | Chr 7 | 264 | 403 | 0.781 | 0.435 | 0.666 |
| Chr 19 | Chr 8 | 264 | 509 | 1.431 | 0.152 | 0.356 |
| Chr 19 | Chr 9 | 264 | 378 | 0.439 | 0.661 | 0.820 |
| Chr 19 | Z1 | 264 | 677 | 3.918 | $8.91*10^{-5}$ | **0.001** |
| Chr 19 | Z2 | 264 | 429 | 6.175 | $6.60*10^{-10}$ | **$3.35*10^{-8}$** |
| Chr 19 | Z3 | 264 | 280 | 3.298 | 0.001 | **0.009** |
| Chr 20 | Chr 21 | 286 | 275 | 1.208 | 0.227 | 0.463 |
| Chr 20 | Chr 22 | 286 | 262 | 3.100 | 0.002 | **0.015** |
| Chr 20 | Chr 23 | 286 | 273 | 1.531 | 0.126 | 0.309 |
| Chr 20 | Chr 24 | 286 | 219 | 0.502 | 0.615 | 0.786 |
| Chr 20 | Chr 25 | 286 | 214 | 3.856 | 0.000 | **0.001** |
| Chr 20 | Chr 26 | 286 | 287 | 1.971 | 0.049 | 0.166 |
| Chr 20 | Chr 27 | 286 | 184 | 1.677 | 0.094 | 0.258 |
| Chr 20 | Chr 28 | 286 | 280 | 3.107 | 0.002 | **0.014** |
| Chr 20 | Chr 29 | 286 | 252 | 1.204 | 0.229 | 0.463 |

| | | | | | | |
|---|---|---|---|---|---|---|
| Chr 20 | Chr 3 | 286 | 572 | 1.985 | 0.047 | 0.162 |
| Chr 20 | Chr 4 | 286 | 492 | 2.303 | 0.021 | 0.089 |
| Chr 20 | Chr 5 | 286 | 544 | 0.991 | 0.322 | 0.553 |
| Chr 20 | Chr 7 | 286 | 403 | 1.529 | 0.126 | 0.309 |
| Chr 20 | Chr 8 | 286 | 509 | 2.231 | 0.026 | 0.103 |
| Chr 20 | Chr 9 | 286 | 378 | 1.168 | 0.243 | 0.478 |
| Chr 20 | Z1 | 286 | 677 | 4.831 | $1.36*10^{-6}$ | **$2.51*10^{-5}$** |
| Chr 20 | Z2 | 286 | 429 | 7.066 | $1.59*10^{-12}$ | **$2.15*10^{-10}$** |
| Chr 20 | Z3 | 286 | 280 | 4.035 | $5.45*10^{-5}$ | **0.001** |
| Chr 21 | Chr 22 | 275 | 262 | 1.889 | 0.059 | 0.188 |
| Chr 21 | Chr 23 | 275 | 273 | 0.322 | 0.747 | 0.872 |
| Chr 21 | Chr 24 | 275 | 219 | -0.628 | 0.530 | 0.729 |
| Chr 21 | Chr 25 | 275 | 214 | 2.705 | 0.007 | **0.040** |
| Chr 21 | Chr 26 | 275 | 287 | 0.743 | 0.457 | 0.683 |
| Chr 21 | Chr 27 | 275 | 184 | 0.593 | 0.553 | 0.736 |
| Chr 21 | Chr 28 | 275 | 280 | 1.876 | 0.061 | 0.193 |
| Chr 21 | Chr 29 | 275 | 252 | 0.023 | 0.981 | 0.996 |
| Chr 21 | Chr 3 | 275 | 572 | 0.569 | 0.570 | 0.753 |
| Chr 21 | Chr 4 | 275 | 492 | 0.920 | 0.358 | 0.590 |
| Chr 21 | Chr 5 | 275 | 544 | -0.400 | 0.689 | 0.835 |
| Chr 21 | Chr 7 | 275 | 403 | 0.207 | 0.836 | 0.922 |
| Chr 21 | Chr 8 | 275 | 509 | 0.840 | 0.401 | 0.634 |
| Chr 21 | Chr 9 | 275 | 378 | -0.132 | 0.895 | 0.954 |
| Chr 21 | Z1 | 275 | 677 | 3.338 | 0.001 | **0.008** |
| Chr 21 | Z2 | 275 | 429 | 5.663 | $1.49*10^{-8}$ | **$4.32*10^{-7}$** |
| Chr 21 | Z3 | 275 | 280 | 2.795 | 0.005 | **0.032** |
| Chr 22 | Chr 23 | 262 | 273 | -1.567 | 0.117 | 0.297 |
| Chr 22 | Chr 24 | 262 | 219 | -2.402 | 0.016 | 0.073 |
| Chr 22 | Chr 25 | 262 | 214 | 0.906 | 0.365 | 0.597 |
| Chr 22 | Chr 26 | 262 | 287 | -1.175 | 0.240 | 0.476 |
| Chr 22 | Chr 27 | 262 | 184 | -1.108 | 0.268 | 0.503 |
| Chr 22 | Chr 28 | 262 | 280 | -0.045 | 0.964 | 0.989 |
| Chr 22 | Chr 29 | 262 | 252 | -1.825 | 0.068 | 0.204 |
| Chr 22 | Chr 3 | 262 | 572 | -1.627 | 0.104 | 0.282 |
| Chr 22 | Chr 4 | 262 | 492 | -1.227 | 0.220 | 0.455 |
| Chr 22 | Chr 5 | 262 | 544 | -2.562 | 0.010 | 0.053 |
| Chr 22 | Chr 7 | 262 | 403 | -1.850 | 0.064 | 0.198 |
| Chr 22 | Chr 8 | 262 | 509 | -1.318 | 0.188 | 0.419 |
| Chr 22 | Chr 9 | 262 | 378 | -2.159 | 0.031 | 0.121 |
| Chr 22 | Z1 | 262 | 677 | 1.039 | 0.299 | 0.532 |
| Chr 22 | Z2 | 262 | 429 | 3.499 | $4.67*10^{-4}$ | **0.005** |
| Chr 22 | Z3 | 262 | 280 | 0.863 | 0.388 | 0.625 |
| Chr 23 | Chr 24 | 273 | 219 | -0.931 | 0.352 | 0.586 |
| Chr 23 | Chr 25 | 273 | 214 | 2.399 | 0.016 | 0.073 |
| Chr 23 | Chr 26 | 273 | 287 | 0.416 | 0.677 | 0.827 |
| Chr 23 | Chr 27 | 273 | 184 | 0.304 | 0.761 | 0.873 |
| Chr 23 | Chr 28 | 273 | 280 | 1.548 | 0.122 | 0.302 |
| Chr 23 | Chr 29 | 273 | 252 | -0.292 | 0.770 | 0.878 |
| Chr 23 | Chr 3 | 273 | 572 | 0.193 | 0.847 | 0.932 |
| Chr 23 | Chr 4 | 273 | 492 | 0.552 | 0.581 | 0.758 |
| Chr 23 | Chr 5 | 273 | 544 | -0.770 | 0.441 | 0.671 |
| Chr 23 | Chr 7 | 273 | 403 | -0.144 | 0.885 | 0.946 |
| Chr 23 | Chr 8 | 273 | 509 | 0.471 | 0.638 | 0.806 |
| Chr 23 | Chr 9 | 273 | 378 | -0.478 | 0.632 | 0.805 |
| Chr 23 | Z1 | 273 | 677 | 2.945 | 0.003 | **0.022** |
| Chr 23 | Z2 | 273 | 429 | 5.294 | $1.19*10^{-7}$ | **$2.85*10^{-6}$** |
| Chr 23 | Z3 | 273 | 280 | 2.466 | 0.014 | 0.064 |
| Chr 24 | Chr 25 | 219 | 214 | 3.157 | 0.002 | **0.013** |
| Chr 24 | Chr 26 | 219 | 287 | 1.333 | 0.183 | 0.412 |
| Chr 24 | Chr 27 | 219 | 184 | 1.134 | 0.257 | 0.492 |

| | | | | | | |
|---|---|---|---|---|---|---|
| Chr 24 | Chr 28 | 219 | 280 | 2.396 | 0.017 | 0.073 |
| Chr 24 | Chr 29 | 219 | 252 | 0.638 | 0.524 | 0.729 |
| Chr 24 | Chr 3 | 219 | 572 | 1.241 | 0.215 | 0.448 |
| Chr 24 | Chr 4 | 219 | 492 | 1.553 | 0.121 | 0.302 |
| Chr 24 | Chr 5 | 219 | 544 | 0.341 | 0.733 | 0.862 |
| Chr 24 | Chr 7 | 219 | 403 | 0.871 | 0.384 | 0.623 |
| Chr 24 | Chr 8 | 219 | 509 | 1.482 | 0.138 | 0.326 |
| Chr 24 | Chr 9 | 219 | 378 | 0.547 | 0.585 | 0.761 |
| Chr 24 | Z1 | 219 | 677 | 3.802 | $1.43*10^{-4}$ | **0.002** |
| Chr 24 | Z2 | 219 | 429 | 5.952 | $2.65*10^{-9}$ | **$1.07*10^{-7}$** |
| Chr 24 | Z3 | 219 | 280 | 3.261 | 0.001 | **0.010** |
| Chr 25 | Chr 26 | 214 | 287 | -2.036 | 0.042 | 0.148 |
| Chr 25 | Chr 27 | 214 | 184 | -1.890 | 0.059 | 0.188 |
| Chr 25 | Chr 28 | 214 | 280 | -0.962 | 0.336 | 0.569 |
| Chr 25 | Chr 29 | 214 | 252 | -2.630 | 0.009 | **0.046** |
| Chr 25 | Chr 3 | 214 | 572 | -2.556 | 0.011 | 0.053 |
| Chr 25 | Chr 4 | 214 | 492 | -2.165 | 0.030 | 0.120 |
| Chr 25 | Chr 5 | 214 | 544 | -3.422 | 0.001 | **0.006** |
| Chr 25 | Chr 7 | 214 | 403 | -2.723 | 0.006 | **0.039** |
| Chr 25 | Chr 8 | 214 | 509 | -2.254 | 0.024 | 0.099 |
| Chr 25 | Chr 9 | 214 | 378 | -3.004 | 0.003 | **0.019** |
| Chr 25 | Z1 | 214 | 677 | -0.100 | 0.920 | 0.965 |
| Chr 25 | Z2 | 214 | 429 | 2.281 | 0.023 | 0.093 |
| Chr 25 | Z3 | 214 | 280 | -0.102 | 0.918 | 0.965 |
| Chr 26 | Chr 27 | 287 | 184 | -0.066 | 0.948 | 0.974 |
| Chr 26 | Chr 28 | 287 | 280 | 1.149 | 0.250 | 0.487 |
| Chr 26 | Chr 29 | 287 | 252 | -0.703 | 0.482 | 0.700 |
| Chr 26 | Chr 3 | 287 | 572 | -0.290 | 0.772 | 0.878 |
| Chr 26 | Chr 4 | 287 | 492 | 0.088 | 0.930 | 0.971 |
| Chr 26 | Chr 5 | 287 | 544 | -1.265 | 0.206 | 0.435 |
| Chr 26 | Chr 7 | 287 | 403 | -0.602 | 0.547 | 0.731 |
| Chr 26 | Chr 8 | 287 | 509 | 0.003 | 0.998 | 0.998 |
| Chr 26 | Chr 9 | 287 | 378 | -0.934 | 0.350 | 0.586 |
| Chr 26 | Z1 | 287 | 677 | 2.499 | 0.012 | 0.060 |
| Chr 26 | Z2 | 287 | 429 | 4.914 | $8.92*10^{-7}$ | **$1.73*10^{-5}$** |
| Chr 26 | Z3 | 287 | 280 | 2.078 | 0.038 | 0.138 |
| Chr 27 | Chr 28 | 184 | 280 | 1.083 | 0.279 | 0.512 |
| Chr 27 | Chr 29 | 184 | 252 | -0.562 | 0.574 | 0.755 |
| Chr 27 | Chr 3 | 184 | 572 | -0.174 | 0.862 | 0.940 |
| Chr 27 | Chr 4 | 184 | 492 | 0.147 | 0.883 | 0.946 |
| Chr 27 | Chr 5 | 184 | 544 | -1.009 | 0.313 | 0.545 |
| Chr 27 | Chr 7 | 184 | 403 | -0.453 | 0.651 | 0.815 |
| Chr 27 | Chr 8 | 184 | 509 | 0.074 | 0.941 | 0.972 |
| Chr 27 | Chr 9 | 184 | 378 | -0.745 | 0.456 | 0.683 |
| Chr 27 | Z1 | 184 | 677 | 2.191 | 0.028 | 0.113 |
| Chr 27 | Z2 | 184 | 429 | 4.323 | $1.54*10^{-5}$ | **$2.50*10^{-4}$** |
| Chr 27 | Z3 | 184 | 280 | 1.905 | 0.057 | 0.185 |
| Chr 28 | Chr 29 | 280 | 252 | -1.810 | 0.070 | 0.207 |
| Chr 28 | Chr 3 | 280 | 572 | -1.611 | 0.107 | 0.286 |
| Chr 28 | Chr 4 | 280 | 492 | -1.202 | 0.229 | 0.463 |
| Chr 28 | Chr 5 | 280 | 544 | -2.567 | 0.010 | 0.053 |
| Chr 28 | Chr 7 | 280 | 403 | -1.838 | 0.066 | 0.202 |
| Chr 28 | Chr 8 | 280 | 509 | -1.295 | 0.195 | 0.426 |
| Chr 28 | Chr 9 | 280 | 378 | -2.152 | 0.031 | 0.121 |
| Chr 28 | Z1 | 280 | 677 | 1.118 | 0.263 | 0.500 |
| Chr 28 | Z2 | 280 | 429 | 3.621 | $2.93*10^{-4}$ | **0.003** |
| Chr 28 | Z3 | 280 | 280 | 0.923 | 0.356 | 0.590 |
| Chr 29 | Chr 3 | 252 | 572 | 0.525 | 0.600 | 0.770 |
| Chr 29 | Chr 4 | 252 | 492 | 0.867 | 0.386 | 0.624 |
| Chr 29 | Chr 5 | 252 | 544 | -0.415 | 0.678 | 0.827 |

| | | | | | | |
|---|---|---|---|---|---|---|
| Chr 29 | Chr 7 | 252 | 403 | 0.177 | 0.860 | 0.940 |
| Chr 29 | Chr 8 | 252 | 509 | 0.790 | 0.430 | 0.666 |
| Chr 29 | Chr 9 | 252 | 378 | -0.154 | 0.878 | 0.946 |
| Chr 29 | Z1 | 252 | 677 | 3.207 | 0.001 | **0.011** |
| Chr 29 | Z2 | 252 | 429 | 5.486 | $4.12*10^{-8}$ | **$1.04*10^{-6}$** |
| Chr 29 | Z3 | 252 | 280 | 2.709 | 0.007 | **0.040** |

**Supplementary table 5.** Results of a linear model to test for a faster Z effect using the formula: log(ω) ~ chromosome type + GC%-4fold + log(FPKM). Significant results are highlighted in bold.

| Coefficient | Estimate | Std. Error | t value | P-value |
|---|---|---|---|---|
| Intercept | -1.556479 | 0.040115 | -38.800 | **$< 2.00*10^{-16}$** |
| anc Z | 0.177458 | 0.061191 | 2.900 | **$3.75*10^{-3}$** |
| neo Z | 0.228779 | 0.042441 | 5.391 | **$7.30*10^{-8}$** |
| gc4fold | -0.806730 | 0.093928 | -8.589 | **$< 2.00*10^{-16}$** |
| log(FPKM) | -0.002042 | 0.006324 | -0.323 | 0.747 |

**Supplementary table 6.** Test statistics and p-values for multiple comparisons of ω between sex biased genes on autosomes and the different Z chromosomes with Kruskal-Wallis rank sum test and pair-wise comparisons with Wilcoxon rank sum test. Bonferroni corrected p-values are shown for all multiple pairwise comparisons. Significant values are highlighted in bold.

| Linkage | Comparison | Test statistic | P-value |
|---|---|---|---|
| Autosomes | FBG vs MBG vs UBG | $\chi^2 = 9.4473$ | **0.009** |
| | FBG vs MBG | $W = 6.36 \times 10^5$ | 0.227 |
| | FBG vs UBG | $W = 3.44 \times 10^6$ | 1 |
| | MBG vs UBG | $W = 3.58 \times 10^6$ | **0.007** |
| Z1 | FBG vs MBG vs UBG | $\chi^2 = 9.7705$ | **0.008** |
| | FBG vs MBG | $W = 3\ 350$ | **0.007** |
| | FBG vs UBG | $W = 8\ 673$ | **0.032** |
| | MBG vs UBG | $W = 2.78*10^4$ | 0.426 |
| Z2 | FBG vs MBG vs UBG | $\chi^2 = 4.7876$ | 0.091 |
| Z3 | FBG vs MBG vs UBG | $\chi^2 = 0.012911$ | 0.993 |

**Supplementary table 7.** Test statistics and p-values for multiple comparisons of ω between autosomes and Z chromosomes for each category of sex bias (FBG=Female biased genes, MBG=Male biased genes, UBG=Unbiased genes) with Kruskal-Wallis rank sum test and pairwise comparisons with Wilcoxon rank sum test. Bonferroni corrected p-values are shown for all multiple pairwise comparisons. Significant values are highlighted in bold.

| Sex-bias | Comparison | Test statistic | P-value |
|---|---|---|---|
| FBG | A vs Z1 vs Z2 vs Z3 | $\chi^2 = 28.053$ | **$3.54*10^{-6}$** |
| | A vs Z1 | $W = 1.24*10^4$ | **0.001** |
| | A vs Z2 | $W = 1.08*10^4$ | **0.002** |

| | | | |
|---|---|---|---|
| | A vs Z3 | $W = 2.05*10^{4}$ | 0.810 |
| MBG | A vs Z1 vs Z2 vs Z3 | $\chi^2 = 14.614$ | **0.002** |
| | A vs Z1 | $W = 8.68*10^{4}$ | 1 |
| | A vs Z2 | $W = 3.76*10^{4}$ | **0.001** |
| | A vs Z3 | $W = 2.44*10^{4}$ | 1 |
| UBG | A vs Z1 vs Z2 vs Z3 | $\chi^2 = 61.719$ | **$2.52*10^{-13}$** |
| | A vs Z1 | $W = 1.04*10^{6}$ | **$3.55*10^{-5}$** |
| | A vs Z2 | $W = 5.96*10^{5}$ | **$4.32*10^{-8}$** |
| | A vs Z3 | $W = 3.98*10^{5}$ | **0.002** |

**Supplementary table 8.** Test statistics and p-values for comparisons of estimates of selection (Direction of Selection (DoS) and Tajima's D) between autosomes (A) and the Z chromosomes (Z1-3) with Kruskal-Wallis rank sum test and pair-wise comparisons with Wilcoxon rank sum test. Bonferroni corrected p-values are shown for all multiple pairwise comparisons. Significant results are highlighted in bold.

| Estimate | DoS | | Tajima's D | |
|---|---|---|---|---|
| | Test statistic | P-value | Test statistic | P-value |
| A vs Z1 vs Z2 vs Z3 | $\chi^2 = 44.461$ | **$1.20*10^{-9}$** | $\chi^2 = 95.725$ | **$< 2.20*10^{-16}$** |
| A vs Z1 | $W = 2.58*10^{6}$ | **$1.62*10^{-4}$** | $W = 4.07*10^{6}$ | **$1.20*10^{-17}$** |
| A vs Z2 | $W = 1.63*10^{6}$ | **$5.42*10^{-7}$** | $W = 2.43*10^{6}$ | **0.016** |
| A vs Z3 | $W = 1.18*10^{6}$ | 1 | $W = 1.60*10^{6}$ | **$3.21*10^{-4}$** |
| Z1 vs Z2 | $W = 1.34*10^{5}$ | 0.846 | $W = 1.44*10^{5}$ | **0.008** |
| Z1 vs Z3 | $W = 9.56*10^{4}$ | **0.033** | $W = 9.49*10^{4}$ | 0.738 |
| Z2 vs Z3 | $W = 6.78*10^{4}$ | **0.001** | $W = 7.02*10^{4}$ | 1 |

**Supplementary table 9.** Test statistics and p-values for multiple comparisons of difference in Direction of Selection (DoS) between sex biased genes on different chromosomes with Kruskal-Wallis rank sum test and pair-wise comparisons with Wilcoxon rank sum test. Bonferroni corrected p-values are shown for all multiple pairwise comparisons. Significant results are highlighted in bold.

| Linkage | Comparison | Test statistic | P-value |
|---|---|---|---|
| Autosomes | FBG vs MBG vs UBG | $\chi^2 = 9.3$ | **0.010** |
| | FBG vs MBG | $W = 6.53*\mathbf{10^{-5}}$ | **0.020** |
| | FBG vs UBG | $W = 3.02*\mathbf{10^{-6}}$ | 1 |
| | MBG vs UBG | $W = 2.95*\mathbf{10^{-6}}$ | **0.016** |
| Z(all) | FBG vs MBG vs UBG | $\chi^2 = 9.6959$ | **0.008** |
| | FBG vs MBG | $W = 1.51*\mathbf{10^{4}}$ | 0.321 |
| | FBG vs UBG | $W = 3.90*\mathbf{10^{4}}$ | 1 |

| | | | |
|---|---|---|---|
| | MBG vs UBG | $W = 9.55*\mathbf{10^4}$ | **0.006** |
| Z1 | FBG vs MBG vs UBG | $\chi^2 = 9.4445$ | **0.009** |
| | FBG vs MBG | $W = 2\ 744$ | 0.088 |
| | FBG vs UBG | $W = 6\ 368$ | 0.891 |
| | MBG vs UBG | $W = 2.35*\mathbf{10^4}$ | **0.018** |
| Z2 | FBG vs MBG vs UBG | $\chi^2 = 6.6962$ | **0.035** |
| | FBG vs MBG | $W = 1\ 635$ | 0.080 |
| | FBG vs UBG | $W = 4\ 434$ | 1 |
| | MBG vs UBG | $W = 9\ 040$ | 0.079 |
| Z3 | FBG vs MBG vs UBG | $\chi^2 = 2.6656$ | 0.264 |

**Supplementary table 10.** Test statistics and p-values for multiple comparisons of difference in Tajima's D between sex biased genes on different chromosomes with Kruskal-Wallis rank sum test and pair-wise comparisons with Wilcoxon rank sum test. Bonferroni corrected p-values are shown for all multiple pairwise comparisons. Significant results are highlighted in bold.

| Linkage | Comparison | Test statistic | P-value |
|---|---|---|---|
| Autosomes | FBG vs MBG vs UBG | $\chi^2 = 0.74387$ | 0.689 |
| Z(all) | FBG vs MBG vs UBG | $\chi^2 = 10.058$ | **0.007** |
| | FBG vs MBG | $W = 1.32*\mathbf{10^4}$ | **0.029** |
| | FBG vs UBG | $W = 4.38*\mathbf{10^4}$ | 0.993 |
| | MBG vs UBG | $W = 1.35*\mathbf{10^5}$ | **0.015** |
| Z1 | FBG vs MBG vs UBG | $\chi^2 = 4.117$ | 0.128 |
| Z2 | FBG vs MBG vs UBG | $\chi^2 = 5.6778$ | 0.058 |
| Z3 | FBG vs MBG vs UBG | $\chi^2 = 5.7329$ | 0.057 |
